# *In vivo* CRISPR screens reveal Serpinb9 and Adam2 as regulators of immune therapy response in lung cancer

**DOI:** 10.1101/2022.03.13.484176

**Authors:** Dzana Dervovic, Edward L.Y. Chen, Ahmad Malik, Somi Afiuni-Zadeh, Jonathan Boucher, Jacob M. Berman, Katie Teng, Arshad Ayyaz, YiQing Lü, Masahiro Narimatsu, Geraldine Mbamalu, Sampath Loganathan, Sebastian Martinez, Ricky Tsai, Jong Bock Lee, Li Zhang, Jeffrey Wrana, Philippe P. Roux, Harland W. Jackson, Daniel Schramek

**Affiliations:** Centre for Molecular and Systems Biology, Lunenfeld-Tanenbaum Research Institute, Mount Sinai Hospital, Toronto, Ontario, Canada; Department of Molecular Genetics, University of Toronto, Toronto, Ontario, Canada; Institute for Research in Immunology and Cancer (IRIC), Université de Montréal, Montreal, Quebec, Canada; Department of Biological Sciences, University of Calgary, Calgary, Alberta, Canada; Department of Otolaryngology, Head and Neck Surgery, McGill University, Montreal, QC, Canada; Toronto General Hospital Research Institute, University Health Network, Toronto, Canada; Departments of Laboratory Medicine and Pathobiology, Immunology, University of Toronto, Toronto, Canada; Department of Pathology and Cell Biology, Faculty of Medicine, Université de Montréal, Montreal, Quebec, Canada

## Abstract

How the genetic landscape of a tumor governs the tumor’s response to immunotherapy remains largely elusive. Here, we established a direct *in vivo* CRISPR/Cas9 gene editing methodology to assess the immune-modulatory capabilities of 573 putative cancer genes associated with altered cytotoxic activity in human cancers. Using *Kras*^G12D^- and *Braf*^V600E^-driven mouse lung cancer models, we identify *Serpinb9* and *Adam2* as our top immune suppressive and immune enhancing genes, respectively. Mechanistically, we show that *Serpinb9* ablation in *Kras*^G12D^- and *Braf*^V600E^-mutant lung tumor cells greatly enhances the efficacy of cytotoxic T-cells *in vitro* and *in vivo*. ADAM2 is a cancer testis antigen broadly expressed in human cancers such as lung adenocarcinoma (13.9%), renal (74.7%), prostate (72.4%), uterine (28.6%) and invasive breast (9.5%) cancer. In our mouse models, we show that *Adam2* expression is induced in *Kras*^G12D^- but not *Braf*^V600E^-driven murine lung tumors and that its expression is further enhanced by immunotherapy. We show that loss of *Adam2* significantly decreases *Kras*^G12D^-lung tumor burden but blocks the efficacy of cytotoxic T-cells. Consistently, Adam2 overexpression dramatically increases tumor growth and enhances immunotherapy efficacy. Mechanistically, we find that *Adam2*’s oncogenic function depends on modulating the tumor immune microenvironment by restraining productive type I and type II interferon responses as well as cytokine signaling, reducing the presentation of tumor-associated antigen, and modulating surface expression of several immunoregulatory receptors within *Kras*-driven lung tumors. Adam2 expression also leads to reduced levels of immune checkpoint inhibitors such as Pd-l1, Lag3, Tigit and Tim3. This reduced exhaustion within the tumor microenvironment may explain why *ex vivo* expanded and adoptively transferred cytotoxic T-cells show enhanced cytotoxic efficacy against Adam2 overexpressing lung tumors. Together, our study highlights the power of integrating cancer genomic with *in vivo* CRISPR/Cas9 screens to uncover how cancer-associated genetic alterations control responses to immunotherapies.

## INTRODUCTION

Lung cancer remains one of the leading causes of cancer-related mortality worldwide, with a 5-year survival rate of <20%^1^. Non-small cell lung cancer (NSCLC) accounts for ~85% of lung cancer and is comprised of lung adenocarcinoma (LUAD), lung squamous cell carcinoma (LUSC) and large cell lung carcinoma (LCLC), among which LUAD is the most prevalent subtype. Exposure to tobacco use is the biggest risk factor and together with other environmental toxins is responsible for the high mutational burden observed in NSCLC. The most frequently mutated genes in LUAD are *TP53^2^* (44%), *KRAS^3^* (33%), *KEAP1^4–6^* (17%), *STK11^7^* (17%), *EGFR^8,9^* (14%), *NF1^10^* (11%), *BRAF^11^* (10%), *PIK3CA^12^* (7%), *MET* (7%)^13–15^. LUAD patients are usually treated with surgery, chemotherapy, radiation therapy, targeted therapy, or a combination of these treatments. Efforts to generate molecular targeted therapies have largely focused on frequently mutated genes and led to the development of tyrosine kinase inhibitors (e.g. *EGFR*, *ALK*, *MET*, *NTRK*) or allele-specific inhibitors (eg. *BRAF*^V600E^, *KRAS*^G12C^)^16–24^. However, prolonged treatment with these targeted therapies often results in the development of acquired drug resistance that limits the duration of their clinical benefit. In addition, the use of these targeted therapies is restricted to a relatively small group of patients whose tumors carry the corresponding genetic alterations.

Immune checkpoint inhibitors blocking the PD-1/PDL1 axis or CTLA4 are new therapeutic approaches for lung cancer patients^25,26^. While immunotherapy and induction of immune responses alone offer modest overall response rates, durable responses for some patients have been observed with combinatorial approaches. For instance, the PACIFIC study shows that adding the anti-PDL1 monoclonal antibody durvalumab to chemotherapy improved progression-free survival (PFS) and overall survival (OS) for patients with NSCLC^27,28^; the CheckMate 816 study confirms a significant increase in pathologic complete response with the addition of the anti-PD1 monoclonal antibody nivolumab to neoadjuvant chemotherapy in resectable lung cancer^29,30^; significantly improved disease free survival is reported in the IMpower010 study that utilizes the anti-PDL1 monoclonal antibody atezolizumab and chemotherapy for patients with resected NSCLC lastly^31^. Lastly, a chimeric antigen receptor (CAR) T cell therapy, targeting several antigens (eg. MSLN, MUC1, NY-ESO-1, GPC3, PSCA, EGFR, ROR1, HER2, PDL1) with limited expression in normal tissues but high and/or specific expression in tumor cells, is presently being tested against NSCLC. The fact that that immune checkpoint blockade yields durable responses in only a small limited percentage of patients (<13%)^33^, underscores the need for deepening our understanding of cancer immune biology. Systematically cataloguing immune-regulatory genes altered in cancer holds the promise of improving treatment decisions and stratifying patients into effective immunotherapies. In addition, elucidating molecular mechanisms that render tumors sensitive or resistant to immunotherapy might identify new targets to enhance existing immunotherapies and improve outcomes in lung cancer.

The advent of functional genetic CRISPR/Cas9 screens has provided a platform for unbiased and systematic identification of genes associated with cancer-intrinsic immune evasion, immunotherapy responses, and therapeutic resistance in tumor cells. While extremely valuable, the majority of CRISPR screens are conducted with T-cells cocultured with cancer cells *in vitro*, which fail to fully recapitulate the complexity of the cellular heterogeneity within tissues^34–36^. CRISPR screens utilizing spheroids or organoids are the next level of *in vitro* systems that better address cellular interactions and mimic *in vivo* conditions. For example, genome-wide CRISPR screen in human NSCLC cell line-based 3D spheroid and xenograft tumors revealed that carboxypeptidase D (CPD) depletion prevents tumor growth in spheroids and *in vivo,* but not in 2D culture^37^. However, *in vitro* spheroids and organoids reiterate *in vivo* architecture only to a certain degree, suffer from low reproducibility, and still fail to fully recapitulate the complexity of a living organism. In contrast, *in vivo* screens performed by allografting of syngeneic mouse cancer cells or by using direct or indirect heterotopic or orthotopic xenografting of patient-derived cells (PDXs), resemble more closely a functional tumor microenvironment (TME). These approaches have led to the identification of novel genes implicated in key immune responses required for cancer cell immune escape and/or resistance to ICB therapy. For instance, *in vivo* genome-wide or epigenetic screens identified novel genes involved in IFNγ signaling and antigen presentation (*PTPN2, ADAR1, APLNR*); suppression of tumor-intrinsic immunogenicity (*SETDB1*); immune-editing of TME (*ASAF1, COP1*); and in direct stress response-induced regulation of PDL1 expression (*EIF5*)^38–44^. Thus, CRISPR-based *in vivo* screens can be leveraged to identify new immune-regulatory genes that might serve as prognostic, diagnostic or potential drug targets in cancer. However, the wounding responses and distorted 3D architecture that inadvertently ensues upon grafting cancer cells, can confound the results of such screens.

To overcome the need for orthotopic transplantation and to assess the function of cancer genes within cells embedded in their native tissue architecture, we developed an autochthonous CRISPR/Cas9 lung cancer screening methodology and combined it with adoptive transfer of cytotoxic T-cells specific to a model tumor antigen. This direct autochthonous screen allows for simultaneous functional interrogation of hundreds putative cancer genes within an intact TME comprised of a functional immune system and endogenous signaling provided by the resident tissue surrounding the target cells. We screened 573 genes that are associated with altered immune activity in human tumors^45^ and report the identification of several known genes such as *Serpinb9,* that plays a role in immune evasion in LUAD as well as other cancers^46,47^. Among novel genes, we unveil the role of a poorly characterized cancer-testis antigen, *Adam2,* in establishing a cold TME by blocking type I and II interferon (IFN-I and IFN-II) and TNFα pathways. However, in the presence of activated exogenous tumor-specific CD8 T-cells, Adam2 expression results in increased tumor clearance by enhanced cytotoxicity of adoptively transferred CD8 T-cells.

## RESULTS

### Direct *in vivo* CRISPR immune screen in lung cancer

To functionally screen genes that determine the immune response to lung cancer, we established an *in vivo* CRISPR loss-of-function screening methodology (**Fig. 1a**). We generated a lentiviral construct that harbors a sgRNA for CRISPR/Cas9-mediated gene editing as well as a Cre-recombinase. For efficient antigen presentation and induction of immune responses by lung epithelial cells, the ovalbumin peptide SIINFEKL (OVA), an experimental tumor-associated antigen (TAA), was cloned into the lentiviral backbone (LV-sgRNA-Cre-OVA). We administer LV-sgRNA-Cre-OVA at postnatal day 2 (P2) to the lungs of Lox-Stop-Lox (LSL)-*Kras*^G12D^ or LSL-*Braf*^V600E^ mice crossed to multicolor LSL-Confetti mice using intranasal instillation. Cre-mediated excision of the LSL-cassettes induced expression of *Kras^G12D^* or *Braf^V600E^* triggered expression of one of four fluorescent proteins encoded in the Confetti reporter cassette (**Fig. 1b**). Histological analysis revealed the induction of hundreds of independent clones and ultimately ~600 *Kras*^G12D^- or *Braf*^V600E^-driven tumors within a single lung (**Supplementary Fig. 1a, b**). For the CRISPR/Cas9 screen, we then generated LSL-*Kras*^G12D^ or LSL-*Braf*^V600E^; LSL-Cas9-GFP; LSL-Luc mice, which concomitantly express a conditional Cas9-GFP transgene for CRISPR/Cas9-mediated gene editing as well as a luciferase transgene for non-invasive measurement of lung tumor volume (hereafter termed *Kras^G12D^;*Cas9 and *Braf*^V600E^;Cas9 mice) (**Fig. 1a**).

**Fig. 1.**
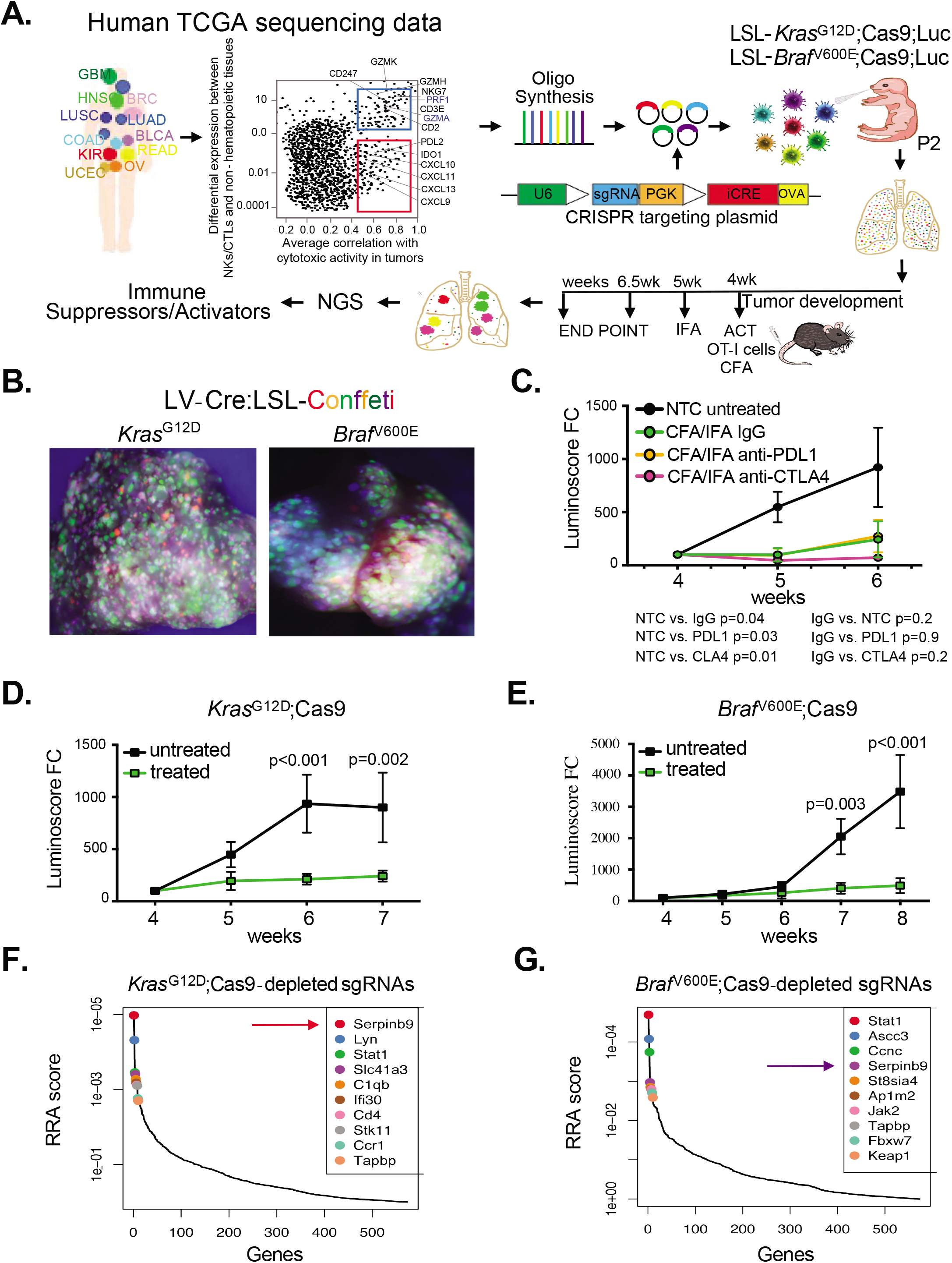
*In vivo* CRISPR screen identifies regulators of immune response in lung cancer. **a**, Schematic showing the workflow of *in vivo* CRISPR screen to identify genes with immune-modulatory cytolytic activity. 573 genes are targeted using a pooled CRISPR sgRNA library in the lung of tumor-prone mice at P2. Mice are treated with OT-I T-cells at week 4, which are activated by CFA/IFA, and sgRNAs in tumor DNA are quantified by NSG. **b**, Representative whole mount immunofluorescence images of the lung from LSL-*Kras*^G12D^;LSL-Confetti and LSL-*Braf*^V600E^;LSL-Confetti mice transduced with Cre lentivirus. **c**, Growth curves of tumors in *Kras*^G12D^;Cas9 mice treated with OT-I cells at week 4 in the presence or absence of PD1 or CTLA-4 blocking antibodies. (n=5 for each group) **d**, Growth curves of tumors in *Kras*^G12D^;Cas9 mice. (untreated, *n*=7; OT-I treated, *n*=10) **e**, Growth curves of tumors in *Braf*^V600E^;Cas9 mice. (untreated, *n*=6; OT-I treated, *n*=6) **f-g**, Top 10 genes whose targeting sgRNAs are depleted in lungs of *Kras*^G12D^;Cas9 and -*Braf* ^V600E^;Cas9 mice. (untreated ,*n*=10; treated, *n*=10) RRA, Robust Rank Aggregation. Data are mean ± s.e.m. Two-sided Student’s t-test.

All LV-sgRNA-Cre-OVA transduced lung cells also express and present the OVA antigen in the context of major histocompatibility complex class I (MHC-I) H-2K^b^ molecules, which can be recognized by adoptively transferred cytotoxic OT-I CD8+ T-cells and can ultimately lead to OT-I-mediated lysis of tumor cells. To optimize different adoptive cell transfer (ACT) immune therapy paradigms, we used OT-I T-cells isolated from OT-I; mT/mG mice that express a membrane-targeted red fluorescent Tomato protein. These cells were injected of *Kras^G12D^*;Cas9 mice via the tail-vain four weeks after lung tumor induction by LV-Cre-OVA inhalation. These mice were then immunized with OVA peptide emulsified in Complete Freund’s adjuvant (CFA) on day 1 and primed with OVA/Incomplete Freund’s adjuvant (IFA) emulsion on day 7 post-ACT to stimulate specific OT-I T-cell responses. We then compared the cytolytic activity of OT-I T-cells isolated from the periphery (spleens and LNs) and lungs of *Kras^G12D^*;Cas9 mice without or with the treatment of PDL-1 or CTLA-4 blocking antibodies by quantifying activation markers and Granzyme (GzmB)-expression. Bioluminescent imaging (BLI) was used to quantify lung tumor burden over time (**Fig. 1c, Supplementary Fig. 2a-c**). Compared to untreated mice, tumor burden in mice treated with OT-I T-cell were significantly smaller, which was further enhanced by CTLA-4 but not PDL1 blocking antibodies. Of note, proliferation and activation of OT-I cells were confirmed *in vitro* and *in vivo* (**Supplementary Fig. 3a-d**). We decided to use the ACT treatment of OVA-CFA/IFA-activated OT-I but without CTLA-4 for the ACT treatment, as this treatment regimen would allow us to find not only genes that block but also genes that enhance the therapeutic effect of ACT.

Next, we generated a pooled lentiviral CRISPR loss-of-function library targeting the mouse homologs of 573 human genes, whose expression, mutation or copy number alteration (CNA) is associated with altered immune cytolytic activity across 8709 tumors from 18 different human cancer types in the TCGA dataset (4 sgRNAs/gene, **Supplementary Table 1**)^45^. We also generated a control library that included 418 non-targeting sgRNAs (termed NTC) as well as an sgRNA targeting the permissive TIGRE locus. These libraries were inhaled into *Kras*^G12D^;Cas9 or *Braf*^V600E^;Cas9 mice (**Fig. 1a**). Next generation sequencing (NGS) confirmed efficient lentiviral transduction of all sgRNAs in the viral library (**Supplementary Fig. 4a, 5a**).

To identify genes that confer resistance or sensitivity to immunotherapy, we quantified sgRNAs at 6.5 weeks after tumor induction, the time point when significant difference in tumor burden was observed between *Kras*^G12D^- or *Braf*^V600E^-driven lung tumors treated and untreated with ACT of activated cytotoxic OT-I T-cells (**Fig. 1d, e**). To reveal suppressors of T-cell mediated killing, we ranked sgRNAs that were depleted in treated versus untreated *Kras*^G12D^;Cas9 or *Braf*^V600E^;Cas9 mice (**Fig. 1f-g, Supplementary Fig. 4 and 5**). *Kras*^G12D^-specific hits included *Serpinb9, Lyn, Stat1, Slc41a* and *C1qb* whereas *Braf*^V600E^-specific hits included *Stat1, Ascc3, Ccnc, Serpinb9* and *St8sia4*. We also observed depletion of sgRNAs targeting genes with known function in tumor immunity and/or resistance to immune checkpoint therapy, such as *CD4*, *C1QB* in *Kras*^G12D^ lungs^48–50^. Depletion of *Stat1*, which has noticeable context-dependent immune-suppressive or -sensitizing role in solid cancers^44,51–53^, was identified in both *Kras*^G12D^ and *Braf*^V600E^ backgrounds. Genes with yet unidentified immunological function in solid cancers included the Src family kinase *Lyn* and the magnesium transporter (*Slc41a3*) identified in the *Kras*^G12D^ screen and transcription coactivator complex (*Ascc3*), haploinsufficient tumor suppressor cyclin C (*Ccnc*) and sialyltransferase (*St8sia4*)^54^ identified in the *Braf*^V600E^ screen (**Fig. 1 f-g**). Together, our direct *in vivo* CRISPR/Cas9 lung cancer screen recapitulated the contribution of previously known immune-regulatory genes and identified potentially new tumor intrinsic factors involved in sensitizing tumor cells to immune-mediated killing.

### Loss of *Serpinb9* increases sensitivity of tumor cells to T-cell mediated killing

Our top hit in the *Kras*^G12D^ background and the fourth hit in the *Braf*^V600E^ background was the serine protease inhibitor (*Serpinb9*). SERPINB9 is the only known intracellular inhibitor of the serine protease granzyme B (GZMB). GZMB is highly expressed by cytolytic CD8^+^ T-cells, natural killer (NK) and γδT-cells to eliminate pathogenic and tumor cells and SERPINB9 is usually expressed in these cytolytic effector cells to protect them from apoptosis induced by their own GZMB^55–57^. In addition, SERPINB9 was recently described as an immunosuppressor and shown to be upregulated in several cancers*^45,46,47^*. For validation experiments, we therefore first focused on *Serpinb9*. To reveal the role of *Serpinb9* in lung cancer *in vivo*, we genetically ablated *Serpinb9* in the lung of *Kras*^G12D^;Cas9 or *Braf*^V600E^;Cas9 mice by inhaling mice with LV-CRE-sg*Serpinb9*-OVA at P2. Three to 4 weeks later, at the time when lung tumors could be detected by BLI, these mice and control littermates transduced with LV-CRE-sgNTC-OVA were injected with OT-I cells followed by OVA-CFA/IFA activation. Efficient depletion of *Serpinb9* was observed in all tested lung tumors (**Supplementary Fig. 6a, b**).

Genetic ablation of *Serpinb9* significantly decreased lung tumor burden in untreated *Kras^G12D^;*Cas9 and *Braf*^V600E^;Cas9 mice (**Fig. 2a, b; Supplementary Fig. 6c-f**). This is in line with the recently identified immunosuppressive role of *Serpinb9* in orthotopic mouse models^47^. While ACT treatment significantly slowed the growth of control *Kras*^G12D^ and *Braf*^V600E^ tumors, loss of *Serpinb9* further enhanced the effect of ACT treatment and completely blocked tumor growth in both *Kras*^G12D^;Cas9 and *Braf*^V600E^;Cas9 mice (**Fig. 2a, b**). Interestingly, although genetic depletion of *Serpinb9* on its own resulted in a similar extension of survival as ACT treatment in control *Kras*^G12D^ mice, it did not further extend the overall survival of ACT-treated *Kras*^G12D^ mice (**Fig. 2c**). This is likely due to the single infusion of OT-I cells at the outset of tumor growth and further optimization would be needed to maximize ACT efficacy. However, in the *Braf*^V600E^-lung cancer model, ACT-treatment significantly increased the overall survival of mice transduced with sg*Serpinb9* compared to control mice (**Fig. 2d**). Together, our findings unveil that depletion of *Serpinb9* enhances CD8 T-cell antitumor immunity in both *Kras*^G12D^- and *Braf*^V600E^-driven mouse models of lung adenocarcinoma.

**Fig. 2.**
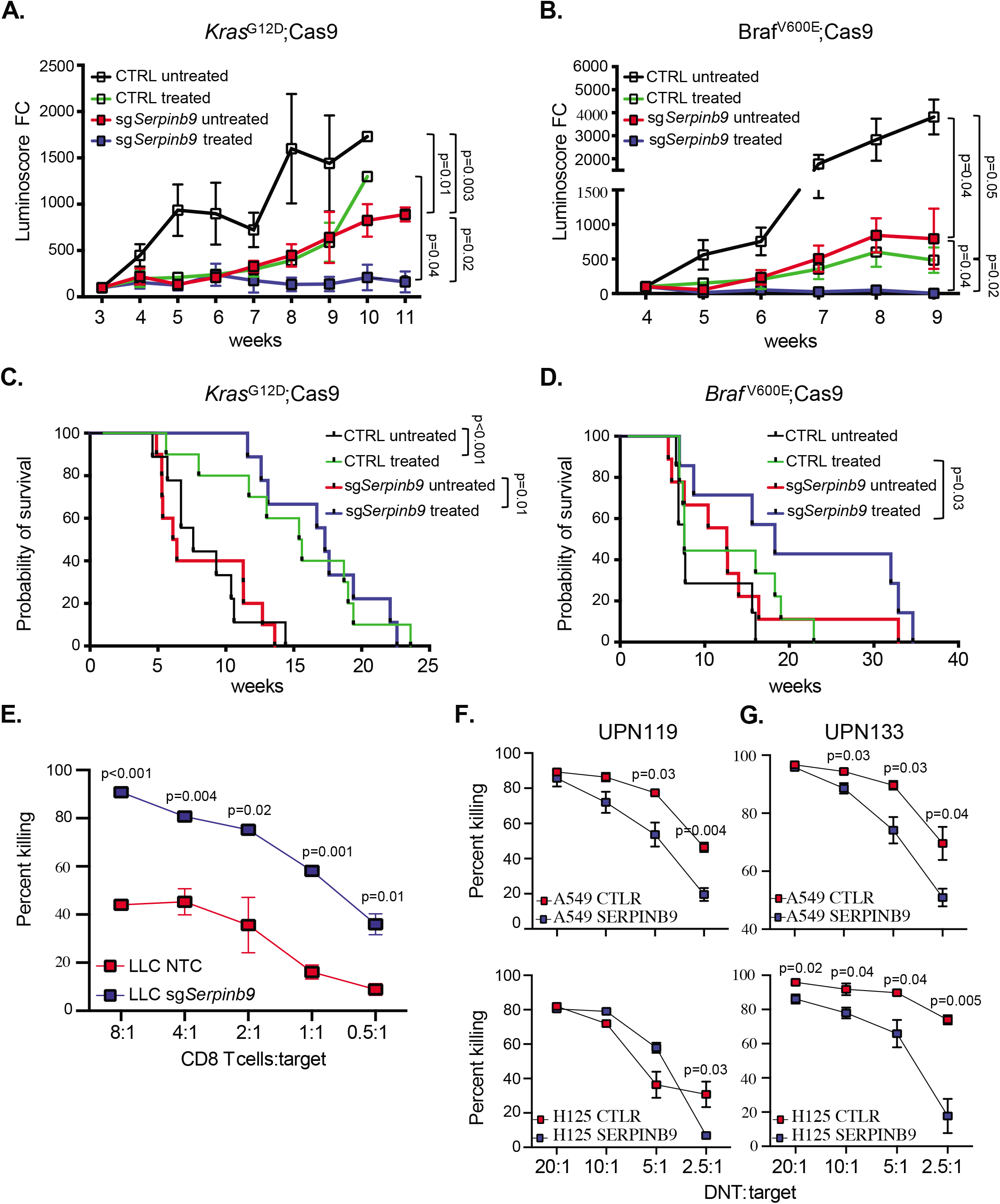
*Serpinb9* deficiency restrains lung tumor growth. **a**, Growth curves of tumors in *Kras*^G12D^;Cas9 mice transduced with sgNTC or sg*Serpinb9* and without (untreated) or with (treated) the ACT treatment. (untreated, *n*=10; treated, *n*=10, for each group) **b**, As in a but in *Braf*^V600E^;Cas9 mice (sgNTC: untreated, *n*=7; treated, *n*=9. sg*Serpinb9*: untreated, *n*=6; treated, *n*=6). **c**, Tumor-free survival of *Kras*^G12D^;Cas9 mice from **(a). d**, Tumor free survival of *Braf* ^V600E^;Cas9 mice in **(b). e**, The effect of *Serpinb9* knockout on T cell-mediated tumor killing. LLC cells transduced with sgNTC or sg*Serpinb9,* and labeled with CFSE were used as targets (T) and cocultured with activated and expended OT-I effector cells (E) at different E:T ratios for 4h. Flow cytometry analysis was used to quantify the percentage of target cells killing at different E:T ratios. Data presents mean ± s.e.m. of three replicates and analysed by two-sided student’s t-test. **f-g**, The effect of SERPINB9 overexpression on DNT cell-mediated killing. Human LUAD cell lines, A549 and H125 labeled with CFSE were used as targets (T) and cocultured with activated and expended DNT effector cells (E) from two different donors (UPN119, UPN133) at different E:T ratios for ~18h. The percentage of target cells killing was determined as in e. Data presents mean ± s.e.m. of three replicates and analysed by two-sided student’s t-test

To further validate the function of *Serpinb9*, we genetically ablated *Serpinb9* in the murine Lewis Lung Carcinoma cells (LLC1), which were established from a C57Bl/6 mouse lung tumor, harbor the activating *Kras^G12C^* mutation and form immunogenic tumors when transplanted into syngeneic mice^58^. Transduction of LLC1 cells with LV-CRE-sg*Serpinb9*-OVA resulted in almost complete depletion of *Serpinb9* (96%) (**Supplementary Fig. 7a**). To assess how Serpinb9 expression modulates cytotoxic T-cell killing, we co-cultured *Serpinb9* knockout LLC1 cells with activated OT-I cells. As expected, *Serpinb9* KO cells were significantly more susceptible to OT-I T-cell mediated killing than control cells (**Fig. 2e, Supplementary Fig. 7b-d**).

Since about 30% of human LUAD have increased *SERPINB9* copy number, which is associated with significantly reduced CD8 T-cell tumor infiltrates and patients’ overall survival, we further validated *SERPINB9* function in human LUAD cell lines (**Supplementary Fig. 8a, b**)^46,59^. We overexpressed human *SERPINB9* in human lung cancer cell lines A549 and H125 and evaluated cytolytic activity of *ex vivo* activated and expanded γδT-cells derived from healthy human donors (UPN119, UPN133), termed DNT cells^60^. As expected, A549 and H125 cell line overexpressing *SERPINB9* were significantly more resistant to human DNT cell-mediated killing when compared to control cells (**Fig. 2f,g; Supplementary Fig. 8c-e, 9 a-d**). Together, our *in vitro and in vivo* functional analysis corroborate the role of *SERPINB9* as an important tumor-intrinsic mechanism in promoting resistance to immunotherapy.

### Loss of *Adam2* impedes T-cell mediated killing

Next, we turned to enriched sgRNAs in our screens, which delineate genes that enhance T-cell mediated killing. Top hits from the *Kras*^G12D^;Cas9 screen included genes known to be required for immune responses, such as *Tapbl*, encoding a component of the antigen processing and presentation machinery; and *Vhl* (von-Hippel-Lindau), a tumor suppressor gene that enhances NK-cell activation in renal cell carcinoma (**Fig. 3a**)^61^. The top hit from the *Braf*^V600E^;Cas9 screen included *Trem2*, a known regulator of immune response with unknown function in tumor cells (**Supplementary Fig. 10a**)^50,62,63^. Interestingly, *Adam2* (Disintegrin and Metalloprotease Domain-Containing Protein2), scored as top gene in the *Kras*^G12D^ but not in the *Braf*^V600E^ screen. *ADAM2* encodes a non-catalytically metalloprotease-like protein, whose expression is usually restricted to the testis and sperm, where it is essential for the sperm-egg fusion^64^. Aberrant expression of *ADAM2* was also reported in some malignancies^65^.

**Fig. 3.**
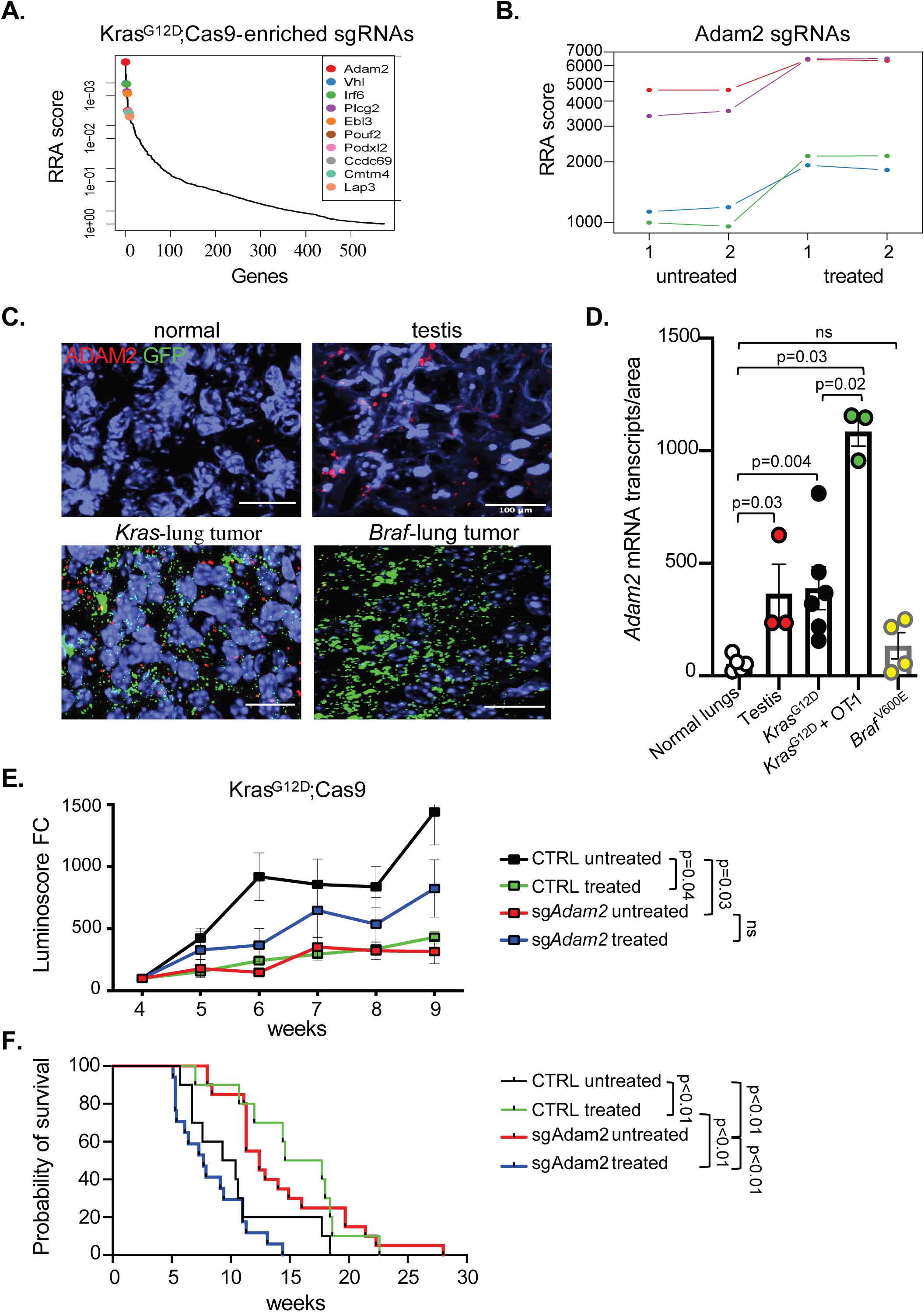
Adam2 is a functional CT antigen in lung cancer. **a**, Top 10 genes whose targeting sgRNAs are in lungs of *Kras*^G12D^;Cas9 screen (untreated, *n*=10; treated, *n*=10). RRA, Robust Rank Aggregation. **b**, sg*Adam2* abundance in untreated lungs and ACT-treated lungs. (n=5 for each replicate) **c**, RNA scope analysis for *Adam*2 (red) and GFP (green; tumor cells) expression in indicated tissues. **d**, Quantification of *Adam*2 transcripts in indicated tissues. (p-values calculated by two-sided student’s t-test; ns = non-significant; data are mean ± s.e.m.) **e**, Growth curves of tumors in *Kras*^G12D^;Cas9 mice transduced with sgNTC (*n*=8, untreated; n=12, treated) or sg*Adam2* (untreated, *n*=14; treated, *n*=17; sg1 and sg2 are shown combined here and are shown separately in SI Fig. 13d, e). **f**, Tumor-free survival of *Kras*^G12D^;Cas9 mice transduced with sgNTC (untreated, n=10; treated, *n=*10) vs. sgRNA sg*Adam2* (untreated, *n=*20; treated, n=18; sg1 and sg2 are shown combined here and are shown separately in SI Fig. 13f, g; data are mean ± s.e.m.)

We first evaluated the expression of *ADAM2* in TCGA LUAD and LUSC samples. Interestingly, 71/510 (13.9%) of LUAD and 24/484 (4.9%) of LUSC tumors showed *ADAM2* expression and ~50% of these cases showed *ADAM2* amplifications, indicating that *ADAM2* amplification correlates with ADAM2 expression (**Supplementary Fig. 10b, c**). In addition, 5% of all lung tumors harbored *ADAM2* missense mutations and 34% possessed gains or amplifications of the genomic region encompassing the *ADAM2* (**Supplementary Fig. 11a-c**). Pan-cancer analysis for *ADAM2* showed a high frequency of expression in breast (9.5%), bladder (16.7%), prostate (72.4%) and renal (74.7%) cancers (**Supplementary Fig. 10b** and **Supplementary Table 2**). Interestingly, amplification of the chromosomal region encompassing *ADAM2* has previously also been associated with low-cytotoxicity in a pan-cancer TCGA analysis^45^. Together, these data show that *ADAM2* is a poorly characterized cancer testis antigen expressed in a wide variety of tumor tissues, including a high proportion of human lung cancers.

To evaluate *Adam2* expression in our mouse lung cancer models, we performed RNA scope analysis. We examined untreated or ACT-treated lung tumors from *Kras*^G12D^;Cas9 or *Braf*^V600E^;Cas9 mice using a probe against *Adam2* as well as a probe against eGFP to detect tumor cells. We used testis as a positive control and normal lungs as a negative control. As expected, we observed high expression of *Adam2* in testis, but not in healthy lungs (**Fig. 3c, d; Supplementary Fig. 12**). Importantly, *Adam2* was expressed in *Kras*^G12D^ lung tumors to a level similar to that seen in testis and its expression was even further increased upon treatment with cytotoxic antigen-specific T-cells. In contrast, *Adam2* mRNA levels in *Braf*^V600E^ lung tumors were negligible (**Fig. 3c, d; Supplementary Fig. 12**). This data corroborates that *Adam*2 is a cancer testis antigen, whose expression is induced in an oncogene-specific manner that can be further increased under selective immune pressure. In addition, the absence of *Adam2* expression in *Braf*^V600E^ tumors likely explains why *Adam2* did not surface as a hit in the *Braf* screen.

To functionally investigate whether *Adam2* regulates anti-tumor immunity, we genetically depleted *Adam2* in lungs of *Kras^G12D^*;Cas9 mice using LV-CRE-sg*Adam2*-OVA. Efficient mutagenesis of *Adam2* was confirmed in all tested lung tumors (**Supplementary Fig. 13a, b**). Next, we evaluated tumor development in untreated versus ACT-treated *Kras^G12D^;*Cas9 mice transduced with sgNTC or 2 independent sgRNAs targeting *Adam2*. We observed that loss of *Adam2* significantly reduced tumor growth and significantly extended the survival of *Kras^G12D^* mice (**Fig. 3e, f** and **Supplementary Fig. 13c-f**). In addition, although ACT-treatment significantly reduced the tumor burden in control lungs, it did not have a significant therapeutic effect in sg*Adam2*-deficient lungs – in fact there was a trend towards increased growth of treated *Adam2*-deficient tumors (**Fig. 3e** and **Supplementary Fig. 13c-g**). Similarly, the overall survival of ACT-treated mice with *Adam2*-knockout tumors was reduced compared to untreated mice with *Adam2* knockout mice, indicating an adverse effect of the ACT-treatment. Collectively, this data shows that *Adam2* functions as a tumor promoter in untreated *Kras*^G12D^ tumors and is required for cytotoxicity of adaptively transferred T-cells.

### *Adam2* suppresses endogenous IFNα/β, IFNγ, and TNFα responses in LUAD *in vitro* and *in vivo*

To further study the function of Adam2 during tumor development and immune regulation, we overexpressed *Adam2* together with the OVA-peptide SIINFEKL in LLC1 cells (=Adam2 O/E cells). Adam2 overexpression was confirmed by western blotting and interferon gamma (IFNγ)-induced presentation of OVA bound to MHC-I H-2K^b^ was confirmed by flow cytometry (**Supplementary Fig. 14a, b**).

We then transplanted *Adam*2 O/E cells or vector only control cells subcutaneously into immunocompetent C57BL/6 (B6) mice. Overexpression of *Adam2* resulted in a dramatically faster tumor growth and significantly reduced survival compared to control cells (**Fig. 4a; Supplementary Fig. 15a-c**). Compared to immunocompetent mice, both Adam2 O/E and control tumors grew faster in immunodeficient <u>N</u>od-<u>S</u>cid-<u>G</u>amma (NSG) mice, and, importantly, at similar rates (**Fig. 4b; Supplementary Fig. 15d, e and h**). Similarly, Adam2 O/E and control tumors did not show significant difference in tumor growth or overall survival in T-cell deficient nude mice (**Fig. 4c; Supplementary Fig. 13f-h**). Since forced expression of *Adam2* had no impact on tumor growth or survival in immunodeficient hosts, these data indicate that Adam2 suppresses endogenous immune responses that restrain the growth of LLC cells.

**Fig. 4.**
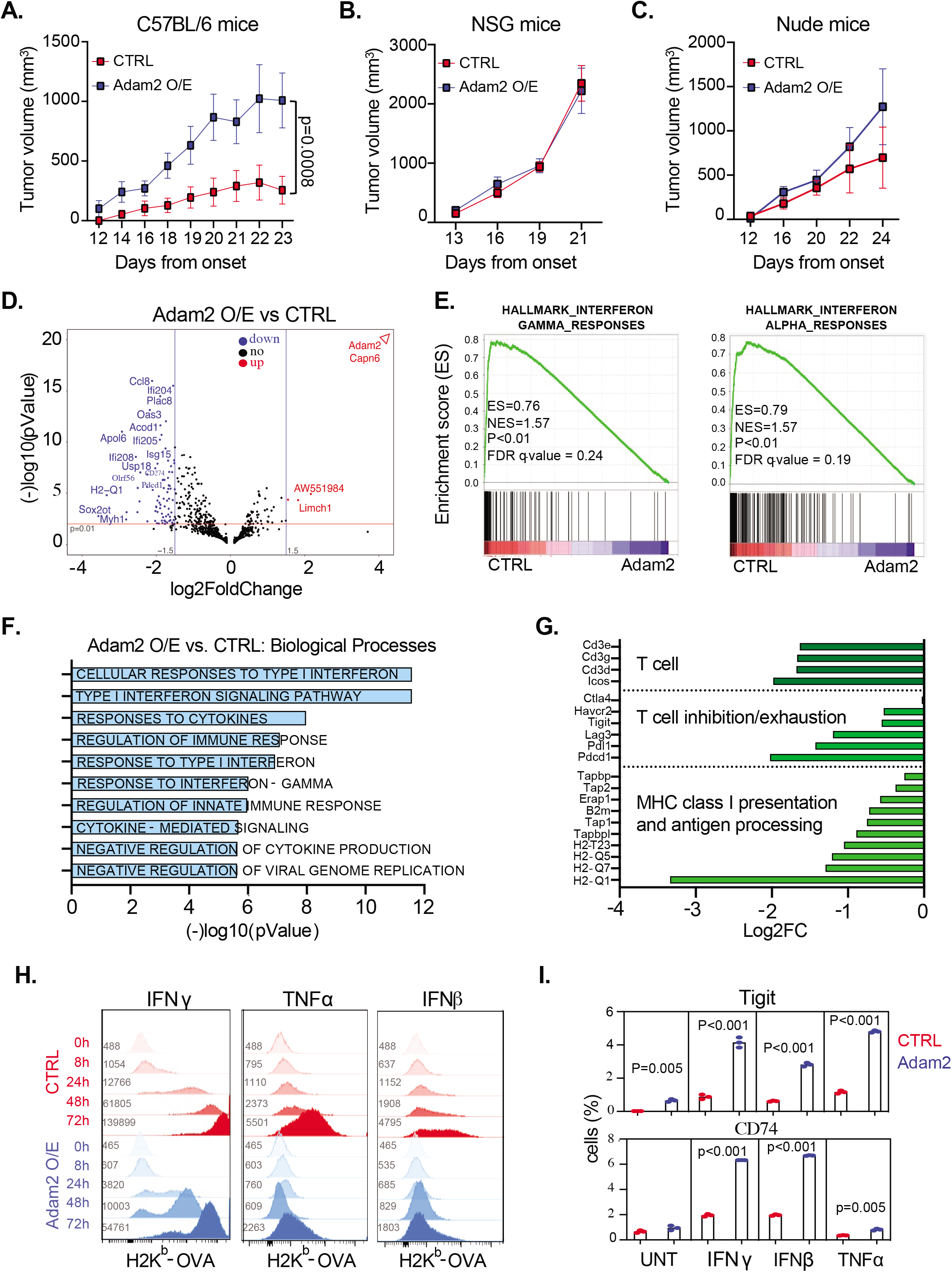
Ectopic expression of *ADAM2* promotes tumorigenesis. **a-c**, Tumor growth of s.c. transplanted CTRL cells or *Adam*2 O/E in B6 mice (n=5) (a); NSG mice (n=5) (b) and Nude mice (n=8) (c). **d**, Volcano plot showing differentially expressed genes between CTLR and Adam2 O/E tumors isolated from B6 mice. **e**, GSEA plots showing Hallmarks of IFNγ and IFNα pathways. **f**, Bar graph showing changes of Gene Ontology of Biological Process between Adam2 O/E versus CTLR tumors. **g**, Expression changes of selected genes from RNA-seq data comparing Adam2 O/E and CTRL tumors. **h**, Cell surface expression of MHC-I H2K^b^-OVA on CTLR and Adam2 O/E cells after treatment with 100ng/ml of IFNγ, IFNβ or TNFα. Grey number denotes mean fluorescence intensity (MFI). i, Percentage of CTLR and Adam2 O/E cells expressing Cd74 and Tigit expression upon IFNγ, IFNβ or TNFα treatment (100ng/ml each, 24h). Data presents mean ± s.e.m. of three replicates and analysed by two-sided student’s t-test.

To elucidate the mechanism through which Adam2 suppresses anti-tumor immunity, we performed whole transcriptome analysis (RNA-seq) on control and Adam2 O/E tumors isolated from B6 mice. Gene set enrichment analysis (GSEA) revealed significant reduction of IFNα/β responses (Cxcl10, Cxcl11, Isg15, Usp18), IFNγ responses (Stat1, Stat2, Tap1, Tapbp, Hla-A, Apol6, Ddx60, Ddhx58), TNFα signaling via NFκβ (Ccl5, Ifih1 Map3k8, Ifit2, Tap1, Il6, Il1b, Irf1), IL2-STAT5 (Icos, Fgl2, Gbp4, Tnfsf10), IL6-JAK-STAT3 (Il18r1, Il2ra, Il1b, Fas) and complement (Ltf, C2, C3, Psmb9, Ccl5, Casp1) pathways (**Fig. 4d-f, Supplementary Fig. 16-20**). In addition, we found downregulation of cluster of differentiation 3 (CD3), inducible T-cell co-stimulator (Icos), several MHC-I molecules, the MHC-I invariant chain B2m, the antigen-processing molecule tapasin (Tap1) as well as downregulation of several checkpoint molecules such as PD1-PDL1 (Pcdc1-CD274), Lag3 (CD223), Tigit and Tim3 (Havcr2) within the tumor microenvironment (**Fig. 4g, Supplementary Fig. 20b**). Thus, multiple cellular responses to INF-I and INF-II, cytokine production and cytokine-mediated signaling as well as negative regulation of overall immune response were suppressed by Adam2.

The prominent role of IFNα/β, IFNγ and TNFα pathways in the induction of antigen processing and MHC-I-mediated antigen presentation on tumor cells prompted us to test H-2K^b^-OVA expression in CTLR and Adam2 O/E cells in response to IFNβ, IFNγ and TNFα treatment^34,35,38,40,66–75^. Flow cytometry revealed markedly delayed and significantly reduced levels of MHCI-OVA surface expression in Adam2 O/E cells in comparison to CTLR cells in response to IFNβ, IFNγ or TNFα stimulation (**Fig. 4h, Supplementary Fig. 21a-g**). Thus, Adam2 overexpression compromises IFNβ, IFNγ or TNFα signaling and MHC-I-mediated antigen presentation blocking, which in turn affect functional innate and adoptive immune responses.

We next tested the expression of other interferon-inducible genes such as PD-L1, CD74, CD44 and TIGIT. Although IFNγ, IFNα and TNFα-induced surface expression of PD-L1 or CD44 was not affected by Adam2, there was a significant increase of CD74 and TIGIT expression in Adam2 O/E cells compared to CTLR cells (**Fig. 4i, Supplementary Fig. 21h-m**). The pro-inflammatory MIF cytokine receptor CD74 is required for antigen presentation by antigen presenting cells (APCs) in the context of MHC class II. However, expression of CD74 is also associated with epithelial cancer cells, where CD74 blocks the MHC-I peptide binding cleft and inhibits TAA presentation to T-cells thus rendering tumors less immunogenic^76, 77^. TIGIT (T cell immunoreceptor with IgG and ITIM domains) was recently identified as a cancer stem cell marker in LUAD^78^ and TIGIT expression on tumor cells was shown to supress CD8 T cells and NK-cells^79^. These results imply that Adam2 downregulates INF-I and INF-II and TNFα responses as well as MHC-I expression, while upregulating other key immune-modulatory receptors resulting in reduced cross-presentation of TAA to antigen-specific CD8 T-cells.

### ADAM2 dictates responses to ACT in LUAD *in vitro* and *in vivo*

Next, we evaluated how Adam2 modulates cytotoxic T-cell killing. *Ex vivo* activated and expanded OT-I CD8^+^-T cells acquired a central memory phenotype, marked by upregulation of activation markers CD25, CD28, CD44, CD62L and GzmB and exhaustion markers CD223 (Lag3) and PD1 (**Supplementary Fig. 22a**). As expected, these activated OT-I cells efficiently killed OVA-peptide expressing CRTL cells. Importantly, overexpression of Adam2 resulted in a significantly enhanced T-cell mediated killing at 4h (p=0.01) and 8h (p=0.0065) (**Fig. 5a; Supplementary Fig. 22b**).

**Fig. 5.**
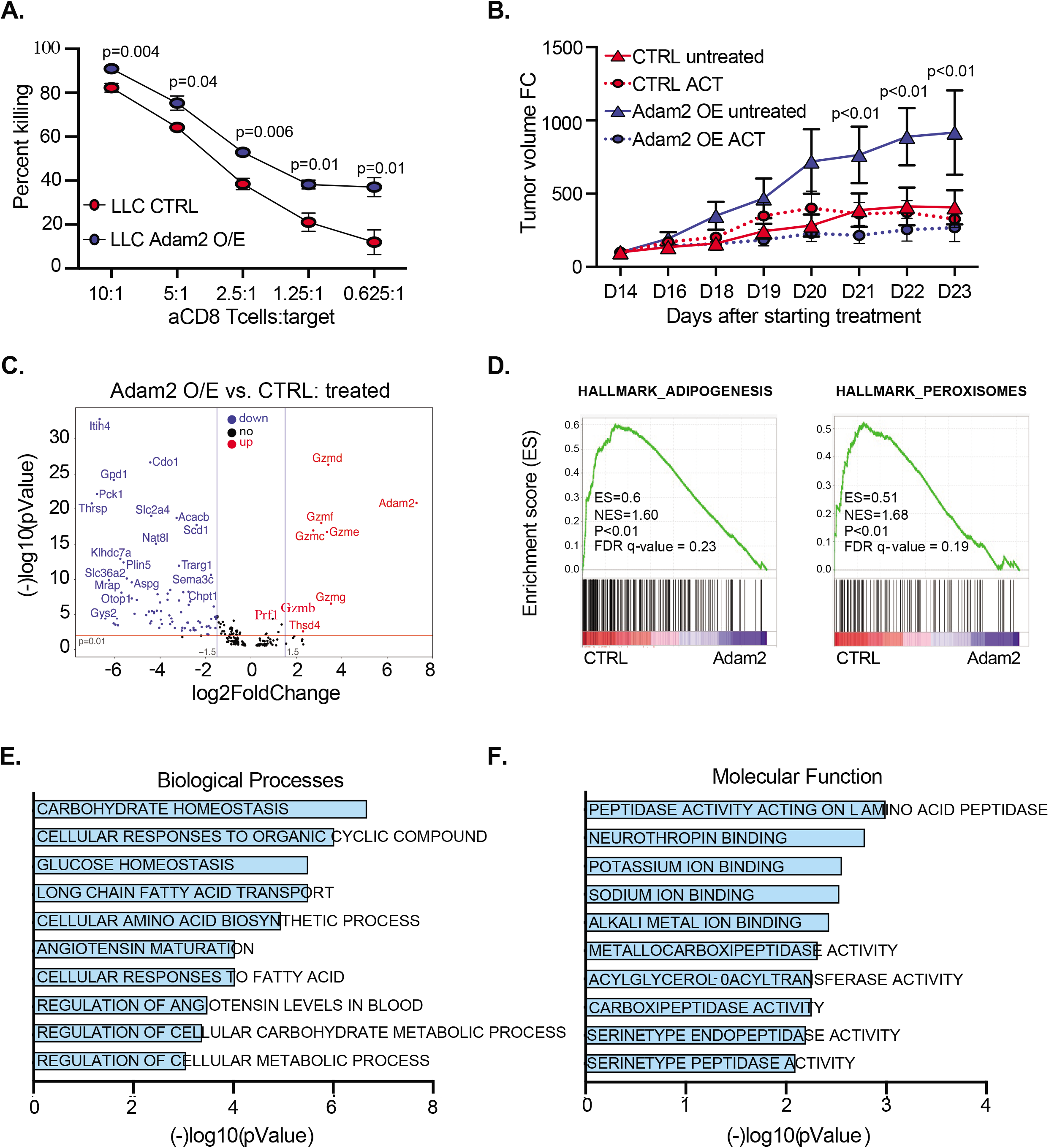
Ectopic expression of *Adam2* invigorates cytolytic activity of TAA-specific CD8 T cells. **a**, The effect of Adam2 overexpression in LLC cells on CD8^+^ OT-I T cell mediated killing. CTRL or Adam2 O/E cells labeled with CFSE were used as targets (T) and cocultured with day 4 activated and expanded OT-I CD8 T-cells (E) at different E:T ratios. Flow cytometry analysis was used to calculate the ratio of killed cells at different E:T ratios. Data presents mean ± s.e.m. of three replicates and analysed by two-sided student’s t-test. **b**, Tumor growth of transplanted CTLR or Adam2 O/E in B6 mice, untreated (n=5) or treated with OT-I T cells (n=5). **c**, Volcano plot showing differentially expressed genes between OT-I treated CTLR and Adam2 O/E tumors isolated from B6 mice. **d**, GSEA plots showing Adipogenesis and Peroxisomes pathways in CTLR tumors versus Adam2 O/E tumors. **e and f**, Bar graph showing changes of Gene Ontology of Biological Process (e) and Molecular Function (f) between OT-I treated CTLR versus Adam2 O/E tumors.

To further understand immunogenic properties of *Adam*2, CTLR or Adam2 O/E cells were transplanted in immunocompetent B6 mice followed by adoptive transfer of OT-I cells at day 14 (i.e., two days after tumor onset). As previously observed, untreated Adam2 O/E tumors grew significantly faster compared to CTLR tumors. Interestingly, activated OT-I had no effect on growth of CTLR tumors, but drastically restrained the growth of Adam2 O/E tumors (**Fig. 5b; Supplementary Fig. 23b**). In addition, transfer of activated OT-I T-cells significantly extended survival of mice bearing Adam2 O/E tumors but not mice bearing CRTL tumors (**Supplementary Fig. 23c**). These results suggest that *Adam2* enhances the cytotoxic potential of antigen-specific CD8 T cells.

Notably, a drastic increase of several serine proteases (GzmB, GzmC, GzmD, GzmE, GzmF, GzmG) was detected by RNA-seq in treated Adam2 O/E tumors compared to treated CTLR tumors isolated from B6 mice (**Fig. 5c; Supplementary Fig. 23d**). In addition, we found upregulation of thrombospondin type1 domain containing 4 (*Thsd4, Adamtsl6*) in treated Adam2 O/E tumors. The expression of *THSD4* is associated with ICB sensitivity in a TCGA pan-cancer analysis^80,81^. Significantly downregulated genes in treated Adam2 O/E tumors were associated with adipogenesis and metabolism with gene sets associated with ‘glucose homeostasis’, ‘cellular responses to fatty acid’, ‘metabolic and amino acid biosynthetic processes’, ‘angiotensin maturation’; and molecular functions associated with ‘peptide activity’, ‘neutrothropin binding’ and ‘carboxypeptidase and serine type peptidase activity’ (**Fig. 5d-f**). While some of downregulated genes have been previously described as targets for cancer therapy such as asparaginase (*Aspg*)^82^, semaphorins (Sema3c)^83^, Adenosin A1 receptor (*Adora*1)^84^, phosphoenolpyruvate (*Pck1*)^85^, their association with immune responses are still largely unexplored. Taken together, these data imply novel functions of Adam2 in the reprogramming of the tumor cells and TME to augment the cytotoxicity of antigen-specific T-cells.

## DISCUSSION

Among the most promising recent advances in oncology is the development of cancer immunotherapies, which largely focus on reactivating endogenous antitumor immune responses. Immune checkpoint inhibitors are designed to break the immune tolerance imposed by the tumor, particularly against cytotoxic T cells^86–90^ and have shown remarkable efficacy against cancers with high mutational burden^91,92^. Similarly, adoptive cell transfer (ACT) of T-cells isolated from patients, genetically modified and activated *in vitro* followed by infusion back into the same patient has proven as a powerful therapeutic strategy^93^. While very effective, immunotherapies result “only” in a ~ 50% response rate and mere 20% of patients experience a durable survival benefit, indicating the existence of resistance mechanisms and other immune escape mechanisms. This raises several questions: What makes certain tumors resistant to immunotherapy and what can we do about it? Are there other immunotherapy targets that we can exploit? Can we combine treatments to enhance the effect of immunotherapies? Are there genetic features that render tumors a priori less or more likely to respond and if so, could we use this information to stratify patients into the right treatment arm? Given the cost of immunotherapies and the burden on health systems, identifying patients who are likely to respond to immunotherapy versus those who will benefit more from other treatments is paramount. From a patient’s perspective, a predictive evaluation of treatment success would be extremely valuable as patients who would unlikely respond to immunotherapy can be evaluated for receiving alternative therapies. As such, cataloguing the genetic changes that determine the sensitivity to immunotherapy would greatly aid clinical decision-making and impact patients’ health.

The explosion of sequencing capabilities is expected to change clinical practices by personalizing treatments based on the genetic make-up of a given tumor and will certainly also help refine immune-oncology. However, one key bottleneck on the path towards ‘Precision ImmunoOncology’ is our fragmented understanding of the functional consequence of most genetic alterations: which mutations are mere bystanders, which are real cancer drivers, and which can be used to determine therapy responses? Determining the effects a given genetic alteration might have on the response to immunotherapy is exceedingly hard and requires a model system that not only ensures a native microenvironment with an intact immune system, but also where tumors can be genetically manipulated and functionally interrogated, ideally in a multiplexed and high-throughput manner. Previous studies have relied mainly on co-culture systems^36,40^ or transplantation models^38^.

To assess how the genomic landscape shapes anti-tumor immunity, we decided to focus on somatic gene alterations that correlate with immune cytolytic activity in the TCGA cohort of 8709 tumors, as spearheaded by Rooney *et al*.^45^. Immune cytolytic activity is based on transcript levels of granzyme A and perforin, two key cytolytic effectors, that are dramatically upregulated upon NK and CD8^+^ T cell activation and during productive clinical responses to immunotherapies^45^. As expected, this approach identified *B2M* (beta-2-microglobulin), *HLA-A*, *-B* and *-C* and *CASP8* (Caspase 8) as significantly mutated genes and *PDL-1/2*, *IDO1/2* as significantly overexpressed genes, highlighting not only loss of antigen presentation and blockade of extrinsic apoptosis as key strategies to overcome cytolytic activity but also validating the approach. In addition to these known immune regulators, Rooney *et al*. identified an additional ~600 significant gene alterations (mutations, CNV, expression) that correlate with cytotoxic index. To functionally assess the tumorigenic and immune-regulatory potential of these genes, we have established a corresponding, multiplexed and barcoded lentiviral CRISPR knock-out library that allowed us to model loss-of-function of these 600 genes directly in the lung of *Kras*^G12D^ and *Braf*^V600E^ mice.

Our top immune-resistance gene is *Serpinb9*. Serpinb*9* directly inhibits GzmB and is highly expressed in cytotoxic T-cell and NK cells to protect those cells from their own GzmB. However, tumor cells hijack this system to protect themselves from cytotoxic attack. Interestingly, Pen Jiang *et al.* identified Serpinb9 as a novel immunosuppressor using a computational method integrating data from the TCGA, PRECOG and METABRIC, which was further substantiated by Liwei Jiang *et al*. showing that Serpinb9 inhibition might constitute a viable therapeutic avenue^46,47^. Here, we show the first autochthonous mouse model of Serpinb9 loss in the lung, corroborate their data, and further extend their findings showing that combining Serpinb9 inhibition with TAA-directed T-cell therapy could yield superior efficacy.

Our top enriched sensitizing gene is *Adam2*. Adam2 is a putative cancer testis antigen, albeit its relevance is still being revealed^65,94^. We found that *ADAM2* is expressed in ~13.9% of human LUAD patients and its expression correlates with *ADAM2* copy number gain or amplification. Interestingly, directly adjacent to *ADAM2* lies *IDO1* and *IDO2* and all three genes are commonly co-amplified in cancer and their amplification is anti-correlated with the cytotoxic index in human tumors^45^. Both, *IDO1* and *IDO2* are known to suppress T-cell immune responses indicating the existence of a strong immune suppressive gene island on chromosome 8p11.21-23.

In mouse tumor models, we could show that *Adam2* expression is induced in *Kras*^G12D^ but not *Braf*^V600E^ tumors and its expression is further increased upon immunotherapy, indicating that *ADAM2* is an oncogene- and immune-responsive *bona fide* cancer testis antigen. In addition, we showed that expression of Adam2 has a dramatic effect on tumor growth by blocking INF-I and INF-II responses as well as several other cytokine signaling pathways thereby restraining the endogenous immune surveillance machinery, affecting predominantly T-cells. To our knowledge, we describe for the first time a cancer testis antigen with not only immunogenic but also with strong immune-modulatory and tumor promoting functions.

Interestingly, we also found that *Adam2* expression not only blocks endogenous cytokine signaling and T-cell responses, but also paradoxically enhances cytotoxicity of adoptively transferred TAA-specific cytotoxic T-cells. One possible explanation is that the reduced endogenous interferon response also reduces expression of interferon-inducible check-point molecules such as PD-L1, Lag3, Tigit and Tim3 in the tumor microenvironment as seen in our mouse model. As such, *Adam2* expression appears to generate a non-exhausted tumor microenvironment, which is highly permissive to *ex vivo* expanded, adoptively transferred cytotoxic T-cells. This could explain why Adam2 overexpressing tumors are actually more sensitive to OT-I cell-mediated killing. Further studies will be needed to elucidate the precise mechanism of how Adam2 regulates interferon and cytokine signaling as well as activation of cytotoxic T cells.

Clinically, our data indicate that patients with tumors expressing ADAM2 would unlikely respond to immune-checkpoint blockade, as this therapy requires a pre-existing productive (albeit somewhat exhausted) endogenous immune response, which are blocked by ADAM2. However, these patients might respond very well to a CAR-T cell therapy or transfer of *ex vivo* expanded and activated autologous TAA-specific T-cells. In addition, inhibiting ADAM2 surface expression or treatment with ADAM2 blocking antibodies (alone or in combination with immune checkpoint inhibitors) might represent new therapeutic avenues for a substantial proportion of cancer patients whose tumors have ectopic ADAM2 expression to rejuvenate their endogenous immune responses to combat their cancers.

Overall, our study highlights how direct *in vivo* CRISPR/Cas9-mediated gene editing can be used to integrate cancer genetics with mouse modeling to elucidate how cancer associated genetic alterations control response to immunotherapies and to identify putative novel immunotherapy targets.

## Data Availability

All RNA-seq data will be made available at NCBI Gene Expression Omnibus.

## Acknowledgements

We thank all members of our laboratories for helpful comments and for their insight and assistance. We also thank The Centre for Phenogenomics, Network Biology Collaborative Centre and Flow Cytometry facility at LTRI as well as the Flow Cytometry Facility at the University of Toronto, BioBox Data Analytics Platform for NGS Data and Histology core at UHN. Funding: This work was supported by a Krembil Foundation Grant, a CCSRI Innovation grant (#706454-1) and a CIHR project grant (#438792) to D.S. and a Terry Fox Research Institute Program Projects Grant to J. W. and D.S et al. (TFRI Project #1107) and a Canadian Cancer Society Impact Grant (grant#: 704121) and a Canadian Institutes of Health Research (grant#: 419699) to LZ. D.D is a recipient of the TD and Medicine by Design Fellowship, S.K.L. is a Canadian Cancer Society Fellowship recipient (BC-F-16#31919). , J.B. is a recipient of the Fonds de recherché du Québec – Santé (FRQS) Fellowship, and P.P.R. is a scholar of the FRQS.

## Author Contributions

D.D. performed all experiments. E.C and HWJ preformed IMC staining. A.M., S.A. and D.D. performed bioinformatic analysis. J.B. and P.P.R. preformed and analysed surfosome proteomics. J.M.B., S.M. and S.L. helped with mouse experiments. K.T. helped with microscopy quantification. A.A. and M.N. and J.W. helped with RNA scope technology. Y.L. helped with cloning and WB analysis. G.M. helped with genotyping and lab work. R.T. helped with qRT-PCR. J-B.L. and L.Z. provided human T cells. D.S. coordinated the project and together with D.D designed the experiments and wrote the manuscript.

## Supplementary Materials

### Supplementary Figure Legends

**Extended data Fig. 1. Histological characterization of *Kras*^G12D^;CAS9 and Braf ^V600E^;Cas9 lung tumors and sgRNA distribution in CRISPR library**

**a-b**, Representative H&E staining of lungs showing hundreds of independent tumors induced by sgNTC by process of intranasal instillation at P2 in *Kras*^G12D^;Cas9 (a) and Braf^V12E^;Cas9 (b) mouse models of LUAD. c, Graph showing sgRNA representation in MEF transduced with plasmid targeting 573 genes=2273 sgRNA pooled CRISPR sgRNA library.

**Extended data Fig. 2. Effects of different treatments on antitumor efficacy after ACT of OT-I CD8^+^ T cells in mice bearing Kras^G12D^ lung-tumor**

**a**, Flow cytometry analysis of CD4 and CD8 expression on total T cells isolated from LNs, spleen and lung of mice subjected to ACT of CD8^+^OT-I cells with IgG2b (*n*=3), anti-PDL1(*n*=4) or anti-CTL4 (*n*=4) treatment (week 5 and 6). **b**, Flow cytometry analysis of CD44 and GzmB expression in endogenous CD8^+^ Tom^−^ and exogenous CD8^+^ Tom^+^ (ACT of OT-I T cells) T cells subjected to different treatments in **a). c-d**, Fold change of total OT-I (c) and OT-I GzmB^+^ **(d)** cell numbers gated on CD8^+^ T cells from mice subjected to different treatments in **a**, was calculated by flow cytometry. Data are mean ± s.e.m. Two-sided Student’s t-test at indicated time-points ns (not significant).

**Extended data Fig. 3. *In vitro* and *in vivo* activation profile of OT-I CD8 T cells**

**a**, Flow cytometry analysis of column purified ,CFSE labeled splenic OT-I CD8 T cells unstimulated or stimulated with anti-CD3/CD28 CD8 T cells for 4 or 7 days. Expression profile of CD25, CD62L selectin, CD44 and GzmB is depicted. CFSE dilution was used to measure T cell proliferation. **b**, Flow cytometry analysis of CD4 and CD8 expression on total T cells isolated from the spleen of B6 untreated mice (group 1, *n*=3) and after ACT of OT-I T cells in B6 mice without treatment (group 2, *n*=3) or with treatment (group 3, *n*=3) with SIINFEKL emulsified in CFA (D1) and IFA (D7). Flow cytometry expression of OT-I TCR-specific chain V⍰2/Vβ5, CD44, CD62L, CD5 and CD25 on gated CD8^+^ T cells. **c**, Total cell number, CD8 T cell number, number of TCR-specific V⍰2/Vβ5^+^ CD8 T cells from group 1, 2 and 3 described in **b** was quantified by flow cytometry. **d**, Total cell number of naïve (CD44^−^CD62L^+^), central memory (CD44^+^CD62L^+^), and activated (CD44^+^CD62L^−^) cells gated on CD8^+^ T cells from group 1, 2 and 3 described in b was quantified by flow cytometry. Results shown are representative of three independent experiments.

**Extended data Fig. 4. CRISPR screens and hits identified after treatment with OT-I cells in *Kras*^G12D^;CAS9 mouse models of LUAD**

**a-c** Graph showing sgRNA representation from two groups of 5 untreated lungs (**a**), two groups of 5 treated lungs after ACT of OT-I cells (**b)** and cumulative representation of sgRNA counts between untreated versus treated lungs (**c**) isolated from *Kras*^G12D^;CAS9 mice transduced with sgRNA library at P2. **d**, Illustration of RRA scores for top depleted/enriched hits from drop-out screen obtained from lungs of *Kras*^G12D^;CAS9 mice transduced with Imm library at P2, before and after treatment.

**Extended data Fig. 5. CRISPR screens and hits identified after treatment with OT-I cells in *Braf*^V600E^;CAS9 mouse models of LUAD**

**a-c** Graph showing sgRNA representation from two groups of 5 untreated lungs (**a**), two groups of 5 treated lungs after ACT of OT-I cells (**b**), and cumulative representation of sgRNA counts between untreated versus treated lungs (**c**) isolated from lungs of *Braf*^V600E^;CAS9 mice transduced with sgRNA library at P2.

**d**, Illustration of RRA scores for top depleted/enriched hits from drop-out screen obtained from lungs of *Braf*^V600E^;CAS9 mice transduced with Imm library at P2, before and after treatment. **e**, Illustration of top 10 enriched genes in lungs of *Braf*^V600E^;CAS9 screen. A comprehensive CRIPSR screen analysis was obtained by use of MAGeCK (<u>M</u>odel-based <u>A</u>nalysis of <u>G</u>enome-wide <u>C</u>RISPR-Cas9 <u>K</u>nockout) algorithm.

**Extended data Fig. 6. SerpinB9 restrains tumor growth in mouse models of lung cancer**

**a-b**, Gene editing efficiency was determined using Sanger sequencing data of PCR-amplified sgRNA target sites followed by <u>T</u>racking of <u>I</u>ndels by <u>D</u>ecomposition (TIDE) algorithm from bulk or GFP sorted tumor cells isolated from untreated or treated lungs of *Kras*^G12D^;CAS9 mice **(a)** or *Braf*^V600E^;CAS9 mice **(b)** transduced with sg*SerpinB9* at P2. **c-d**, Representative images of tumor development in lungs of untreated or treated *Kras*^G12D^;CAS9 mice **(c)** and *Braf*^V600E^;CAS9 mice **(d)** inhaled at P2 with sgNTC or sg*Serpinb9* measured *in vivo* by BLI, weekly. The pseudocolor images (rainbow color scale) were adjusted to the same threshold and the signals are expressed in BLI counts **(e-f)** Grouped bar graphs show luminoscore in radiance (total flux p/s) and are expressed as fold induction from lungs of untreated or treated *Kras*^G12D^;CAS9 mice **(c)** and *Braf*^V600E^;CAS9 mice **(d)** inhaled at P2 with sgNTC or sg*Serpinb9* measured *in vivo* by BLI, weekly. Fold induction of the mean bioluminescence activity represents the mean measured at different time points divided by the mean at the time point (week 3 or 4) when mice were treated with CD8^+^ Tom^+^ cells (baseline). Significance in percent changes in the tumor volume between different groups were calculated by two-sided Student’s t-test at indicated time-points ns (not significant), *P≤0.05, **P≤0.01 and ***P≤0.001. Data are mean ± s.e.m.

**Extended data Fig. 7. Genetic depletion of *Serpinb9* enhances lung tumor susceptibility to T cell-mediated killing *in vitro***

**a**, Gene editing efficiency of *Serpinb9* KO stable murine Lewis Lung Carcinoma (LLC) cell line subclone was determined using Sanger sequencing data of PCR-amplified sgRNA target sites followed by <u>I</u>nference of <u>C</u>RISPR <u>E</u>dits (ICE) analysis. **b**, LLC transduced with sgNTC or sg sg*Serpinb9* labeled with CFSE were used as targets (T) and cocultured with activated and expended OT-I effector cells (E) at different E:T ratio for 4h. Bar graphs show percentage of T cell mediated killing at different E:T transduced with sgNTC (blue) or sg*Serpinb9* (red). Flow cytometry analysis was used to calculate the ratio of PI^−^ CFSE^+^ depleted cells cultured with effector cells and divided with PI^−^ CFSE^+^ depleted cells without effector cells at different time points. Numbers represent fold change of the mean values obtained from triplicate cocultures. **c**, Representative flow cytometry-based killing assay (overnight) of CFSE labeled LLC cells transduced with sgNTC or sg*Serpinb9* (target) by *in vitro* activated OT-I effector cells (E) at different ratios. **d**, Bar graphs represent percentage of T cell-mediated killing of LLC targets transduced with sgNTC (blue) or sg*Serpinb9* (red) over night. Flow cytometry analysis was used to calculate the ratio of PI^−^ CFSE^+^ depleted cells described in **(c)**. Numbers represent fold change of the mean values obtained from triplicate cocultures described in **(c)**.

**Extended data Fig. 8. Expression of *SerpinB9* in human LUAD**

**a**, TCGA analysis depicted as a cBioPortal OncoPrint reveals that 38% (*n*=191) of LUAD (*n*=507) and 27% (*n*=127) of LUSC (*n*=499) have gain or amplification of S*ERPINB*9, together account for 30% of these two subtypes of NSCLC examined. **b**, Tumor Immune Disfunction and Exclusion (TIDE) analysis from TCGA (*n*=487) and GSE14184 (*n*=71) LUAD cohorts illustrates that low expression of SERPINB9 significantly correlates with higher level of CTLs in tumors and overall survival. **c**, Western blot analysis (blotted for V5-tag) showing overexpression of SERPINB9 in A549 and H125 human cancer cell lines. **d**, Representative images of SERPINB9 staining in human lung cancer cell lines A549 and H125 without or with overexpression of SERPINB9. **e**, Quantification of SERPINB9 staining in A549 and H125 cancer cell lines without or with overexpression of SERPINB9.

**Extended data Fig. 9. Overexpression of SerpinB9 in human LUAD cell lines restrains T cell-mediated killing**

**a-d**, The effect of SERPINB9 overexpression on DNT cell-mediated killing was measured by flow cytometry. Human LUAD cell lines A549 and H125 without (blue bars) or with (red bars) SERPINB9 overexpression labeled with CFSE were used as targets (T) and cocultured with day 15 activated and expended DNT cells (E) from two different donors (UPN119, UPN133) at different E:T ratio for ~18h. Flow cytometry analysis was used to calculate the ratio of CFSE^+^ depleted cells cultured with effector cells and divided with CFSE^+^ depleted cells without effector cells at different time points. Numbers represent fold change of the mean values obtained from triplicate cocultures. The difference between overexpression and control conditions were measured by two-sided student’s t-test at indicated time-points.

**Extended data Fig. 10. Pan-cancer expression of Adam2**

**a**, Illustration of top 10 enriched genes in lungs of *Braf*^V600E^;CAS9 screen. A comprehensive CRIPSR screen analysis was obtained by use of MAGeCK (<u>M</u>odel-based <u>A</u>nalysis of <u>G</u>enome-wide <u>C</u>RISPR-Cas9 <u>K</u>nockout) algorithm. **b**, Pan-cancer analysis of Adam2 mRNA expression from TCGA data reveals increased levels of *ADAM*2 transcripts across multiple cancers with the highest expression detected in Bladder Urothelial Carcinoma, Breast Invasive Ductal Carcinoma, Lung Adenocarcinoma, Lung Squamous Carcinoma, Uterine Cancer, Prostate Adenocarcinoma and Chromophobe Renal Carcinoma. c, Bar plot shows alteration (mutation, fusion, amplification, deep deletion, and multiple alterations) frequencies of *ADAM*2 across 33 tumor types obtained from TCGA dataset. *ADAM2* is amplified in ~5% of NSCLC cases analyzed.

**Extended data Fig. 11. Expression of *ADAM2* in human LUAD**

**a,b** TCAG analysis depicted as a cBioPortal OncoPrint reveals that 4% (*n*=20) of LUAD (*n*=507) and 6% (*n*=29) of LUSC (*n*=469) have *ADAM2* missense mutations, together account for 5% of these two subtypes of NSCLC examined (a) and that 28% (*n*=141) of LUAD (*n*=507) and 45% (*n*=210) of LUSC (*n*=469) have gain of *ADAM2*, together account for 34% of these two subtypes of NSCLC examined (b). c, Cancer distribution and CNV distribution of LUAD cases obtained from TCGA data.

**Extended data Fig. 12 Oncogene-induced expression of Adam2 is detected in lung cancer**

Representative 3D images from normal lungs (negative control), testis (positive control), *Kras*^G12D^ and *Braf*^V600E^ lungs inhaled at P2 with NTC virus probed for Adam2 (red), GFP (green) using RNAscope and images were captured with Nikon eclipse Ti2 inverted fluorescent microscope.

**Extended data Fig. 13 *Adam2* deficiency inhibits T cell-mediated immune responses in mouse model of lung cancer**

**a-b**, Gene editing efficiency was determined using Sanger sequencing data of PCR-amplified sgRNA target sites followed by <u>T</u>racking of <u>I</u>ndels by <u>D</u>ecomposition (TIDE) algorithm on bulk or GFP sorted tumor cells isolated from untreated or treated lungs of -*Kras*^G12D^;CAS9 mice transduced with sg1*Adam2* **(a)** or sg2*Adam2* **(b)** at P2. **c**, Representative images of tumor development in lungs of untreated or treated *Kras*^G12D^;CAS9 mice inhaled at P2 with sgNTC or sg*Adam2* measured *in vivo* by BLI, weekly. The pseudocolor images (rainbow color scale) were adjusted to the same threshold and the signals are expressed in BLI counts **(d-e)** Grouped bar graphs show luminoscore in radiance (total flux p/s) and are expressed as fold induction, from untreated or treated lungs of -*Kras*^G12D^;CAS9 mice transduced with sg1*Adam2* (a) or sg2*Adam2* (b). Fold induction of the mean bioluminescence activity represents the mean measured at different time points divided by the mean at the time point (week 3 or 4) when mice were treated with CD8^+^ Tom^+^ cells (baseline). Significance in percent changes in the tumor volume between different groups were calculated by two-sided Student’s t-test at indicated time-points ns (not significant). Data are mean ± s.e.m. **f**, Tumor free survival of *Kras*^G12D^;CAS9 mice transduced with sgNTC (untreated n=10; treated *n=*10) vs. sg1*Adam2* (untreated *n=*8; treated n=10). **g**, Tumor free survival of *Kras*^G12D^;CAS9 mice transduced with sgNTC (untreated n=7; treated *n=*10) vs. sg2*Adam2* (untreated *n=*12; treated n=8). Data are mean ± s.e.m.

**Extended data Fig. 14. Ectopic expression of ADAM2 in LLC cells**

**a**, Western blot analysis showing lack of or expression of ADAM2 in LLC cell line blotted for V5 tag and GAPDH. **b**, Flow cytometry analysis of H2K^b^ or H2K^b^SIINFEKL in LLC cell lines without or with overexpression of ADAM2.

**Extended data Fig. 15. Fig. 4. Ectopic expression of ADAM2 promotes tumorigenesis in the absence of Ag-specific immunity**

**a**, Western blot analysis showing lack of or expression of ADAM2 blotted for V5 tag and GAPDH in tumors isolated from s.c. transplanted CTLR or Adam2 O/E cells (0.1×10^6^) in B6 mice, untreated (n=5) or treated with OT-I cells (n=5). **b**, Fold change of the growth of CTLR or ADAM2 O/E tumors (n=8 for each group) was measured by caliper. **c**, Kaplan-Meier survival analysis of the B6 mice bearing CTLR or ADAM2 O/E tumors, untreated or treated with ACT of OT-I cells. **d-e**, Tumor volume of s.c. transplanted CTLR or Adam2 O/E cells (0.1×10^6^) in NSG (n=8) or Balb/c Nude (n=5) mice for each group was measured by caliper from the day of tumor onset. **f-g**, Western blot analysis showing lack of or expression of ADAM2 blotted for V5 tag and GAPDH in tumors isolated from (d) (f) and from (e) (g). **h**, Kaplan-Meier survival analysis of the host animals (NSG and Nude) bearing CTLR or ADAM2 O/E tumors. The difference between overexpression and control conditions were measured by two-sided student’s t-test at indicated time-points. ns (non-significant), *P≤0.05, **P≤0.01 and ***P≤0.001. Data are mean ± s.e.m.

**Extended data Fig. 16. Adam2 downregulates multiple pathways associated with immune function**

**a-d** Gene set enrichment analysis reveals downregulation of TNFα signaling via NFκβ (**a)**, IL2_STAT5_signaling (**b)**, IL6_JAK_STAT3 signaling (**c)**, and complement pathways (d) in tumors overexpressing ADAM2 in comparison to CTLR tumors.

**Extended data Fig. 17. a, b,** Heat map from GSEA Hallmark genes (a) and Reactome genes (b) of IFNγ signaling in ADAM2 O/E tumors versus CTLR tumors.

**Extended data Fig. 18. a, b,** Heat map from GSEA Hallmark IFNα signaling (a) and Hallmark of TNFα signaling (b) in ADAM2 O/E tumors versus CTLR tumors.

**Extended data Fig. 19. A, b,** Heat map from GSEA Hallmark IL2_STAT5 signaling (a) and Hallmark of IL6_JAK_STAT3 signaling (b) in ADAM2 O/E tumors versus CTLR tumors.

**Extended data Fig. 20. a,** Heat map from GSEA Hallmark Complement in ADAM2 O/E tumors versus CTLR tumors. **b,** (−)Log10 fold changes of Reactome created from the gene ontology tree of biological processes in CTLR versus ADAM2 O/E tumors

**Extended data Fig. 21. Ectopic expression of ADAM2 regulates IFN associated genes**

**a**, Flow cytometry analysis of cell surface expression for H2K^b^/ H2K^b^SIINFEKL on CTLR versus ADAM2 O/E cells untreated or treated with IFNγ, IFNβ or TNFα at 8h, 24h, 48h and 72h. **b-d**, The percentage of H2K^b^/ H2K^b^SIINFEKL cell surface expression on CTLR versus ADAM2 LLCs from groups treated with only IFNγ (**b)**, only IFNβ (**c)**, or only TNFα (**d)**. **e-g**, Total cell numbers of H2K^b^/ H2K^b^SIINFEKL cell surface expression on CTLR versus ADAM2 LLCs from group treated with only IFNγ **(e)**, only IFNβ **(f)**, or TNFα **(g). h**, MFI from flow cytometry analysis for cell surface expression of PDL1 on CTLR versus ADAM2 O/E cells untreated or treated with IFNγ, IFNβ or TNFα at 24h. **i**, percentages from group analyzed in **(h). j**. Total cell numbers from group in **(h). k**, MFI from flow cytometry analysis for cell surface expression of CD74 and Tigit on CTLR versus ADAM2 O/E cells untreated or treated with IFNγ, IFNβ or TNFα at 24h. **l-m**, Total cell numbers from flow cytometry analysis for cell surface expression for CD74 and Tigit on CTLR versus ADAM2 O/E cells untreated or treated with IFNγ, IFNβ or TNFα at 24h.

**Extended data Fig. 22. Overexpression of ADAM2 in the LLC Cell Line augments Ag-specific T-cell mediated killing**

**a**, Flow cytometry analysis of SIINFEKL-activated and expended OT-I splenocytes for 4 days showing cell surface expression of CD8, Tom; activation markers CD25 (IL2R⍰), CD28, CD44, CD62L; exhaustion markers PD1, CD223 (Lag3), Tim3, CTLA4 and cytotoxic marker GzmB versus isotype control. Isotype control is shown in blue and stains in red, where applicable. **b,** The effect of Adam2 overexpression on OT-I cell-mediated killing was measured by flow cytometry. CTLR or Adam2 O/E cells labeled with CFSE were used as targets (T) and cocultured with OT-I cells (E) at different E:T ratio for 8h. The difference between overexpression and control conditions were measured by two-sided student’s t-test.

**Extended data Fig. 23. Overexpression of ADAM2 in the LLC Cell Line augments Ag-specific T-cell mediated killing**

**a**, Western blot analysis showing the lack of or expression of ADAM2 blotted for V5 tag and GAPDH in tumors isolated from s.c. transplanted CTLR or Adam2 O/E cells (0.1×10^6^) in B6 mice, untreated (n=5) or treated with OT-I cells (n=5). **b,** Fold change of tumor growth of s.c. transplanted CTLR or Adam2 O/E cells (0.1×10^6^) in B6 mice, untreated (n=5) or treated with OT-I cells (n=5) for each group was measured by caliper. **c,** Kaplan-Meier survival analysis of B6 mice bearing CTLR or Adam2 O/E tumors, untreated or treated with ACT of OT-I cells. The difference between overexpression and control conditions were measured by two-sided student’s t-test. **d,** Bar graph shows fold change of top 12 enriched genes obtained from RNA-seq data performed on ADAM2 O/E tumors in comparison to CTLR tumors treated with OT-I cells.

## Materials and Methods

### Animals

Equal numbers of male and female animals were used throughout the study without any bias. Animal husbandry, ethical handling of mice and all animal work were carried out according to guidelines approved by Canadian Council of Animal Care and under protocols approved by the Centre for Phenogenomics Animal Care Committee (18-0272H). The animals used in this study, LSL-Kras^G12D^ (008179), LSL-Braf^V600E^ (017837), R26-LSL-CAS9-GFP (026175), FVB.129S6 (B6) *Gt (ROSA) 26Sor^tm1(Luc)Kael^*/J (005125), C57BL/6-Tg(TcraTcrb)1100Mjb/J (003831, also known as OT-I), Gt(ROSA)26Sor^tm4(ACTB-tdTomato,-EGFP)Luo^/J (007576, also known as mT/mG), Gt(ROSA)26Sor^tm1(CAG-Brainbow2.1)Cle^/J (013731, also known as R26R-Confetti), NOD.Cg-Prkdc^scid^ IL2rg^tm1Wjl^/Sz/J (005557, also known as NSG), NU/J (002019, known as Nude) and C57BL/6J (000664) were purchased from the Jackson Laboratory. CRISPR screen in the LSL-Kras ^G12D^-CAS9-LUC and LSL-Braf ^V600E^-CAS9-LUC was performed in a F1 FVBNxC57BL/6J background. Genotyping was performed by PCR using genomic DNA prepared from mouse ear punches.

### Cell lines

A549 and H125 cell lines were purchased from ATCC. LLC cancer cell line was received as a gift from Dr. Hansen He (The Princess Margaret Cancer Centre). All cell lines were tested for mycoplasma and cultured in DMEM (Gibco) media supplemented with 10% fetal bovine serum (FBS) and antibiotics. For some experiment, puromycine or blasticidine selection at 10ug/ml was used.

### Lentiviral constructs and library construction

pLKO sgRNA-Cre plasmid^95^ was modified to express OVA peptide SIINFEKL (gBlock gene fragments, IDT technologies), hereafter pLKO-sgRNA-Cre-P2A-OVA. sgRNAs targeting mouse homologues of human genes associated with immune cytolytic activity, selected from Rooney et al.^45^, were obtained from Hart et al^96^ (4 sgRNAs/gene) and non-targeting sgRNAs were obtained from Sanjana et al.^97^, ordered as a pooled oligo chip (CustomArray Inc., USA) and cloned into sgRNA-Cre-P2A-OVA using BsmBI restriction sites. For validation of top enriched hits, individual sgRNA 1 and 2 targeting *SerpinB9* and *Adam2* were ordered from Sigma and cloned into the same sgRNA-CRE-P2A-OVA plasmid backbone using BsmBI site. We excluded frequent and known immune checkpoint inhibitors such as *CTLA4*, *PDCD1(PD1)*, *PDL1* from the immune genes library. The non-targeting sgRNAs as well as an sgRNA (actgccataacacctaactt) targeting the permissive TIGRE locus^98^ were designed not to target in the mouse genome as negative control.

For overexpression analysis, V5 tagged mouse Adam2, human ADAM2, and human SERPINB9 were obtained from the ORFeome collaboration provided by Fritz Roth, originally from CCSB, DFCI, Harvard. Codon-optimized mouse SerpinB9 was ordered as gBlock fragment from Twist Bioscience. These were cloned into original PLX306 (kindly provided by David Root; Adgene #41391) or modified to express OVA peptide SIINFEKL (PLX306-puromycin-P2A-OVA) construct using gataway cloning system. For *in vivo* experiments puromycin was replaced by iCRE.

### Lentivirus production and transduction

Large-scale production and concentration of lentivirus were performed as previously described^99–102^. Briefly, 293T cells (Invitrogen R700-07) were seeded on a poly-L-lysine coated 15 cm plates and transfected using PEI (polyethyleneimine) method in a non-serum media with lentiviral construct of interest along with lentiviral packaging plasmids psPAX2 and pPMD2.G (Addgene plasmid 12259 and 12260). 8-12 hours post-transfection media was added to the plates supplemented with 10% Fetal Bovine Serum (FBS) and 1% Penicillin-Streptomycin antibiotic solution (w/v). 48 hours later, the viral supernatant was collected and filtered through a Stericup-HV PVDF 0.45-μm filter, and then concentrated ∼2,000-fold by ultracentrifugation in an MLS-50 rotor (Beckman Coulter). Viral titers were determined by infecting the R26-LSL-tdTomato MEFs and FACS based quantification. *In vivo* viral transduction efficiency was determined by injecting decreasing amounts of a single viral aliquot of known titer, diluted to a constant volume of 2x and 10x per intranasal instillation at P2. The percent of infection was analyzed by BLI measuring the total flux [p/s] in lung. 10ul of 1-1.5×10^8^ PFU at P2 was determined as optimal concentration for efficient induction of lung tumorigenesis by week 3-4 in Kras and Braf LUAD mouse models.

### Deep Sequencing: sample preparation, pre-amplification, and sequence processing

Genomic DNA from epithelial and tumor cells were isolated with the DNeasy Blood & Tissue Kit (Qiagen). Genomic DNA concentration was quantified using Qubit dsDNA BR Assay (cat no. Q32853). 40μg genomic DNA of each lung tumor (n=20) was used as template in a pre-amplification reaction by nested primers v2.1-F1 gagggcctatttcccatgattc and v2.1-R1 gttgcgaaaaagaacgttcacgg with 25 cycles and Q5 High-Fidelity DNA Polymerase (NEB), followed by unique barcoded primer combination for pool of all individual 50ul reactions for each genomic DNA sample. 5ul of PCR1 product was run on a 1% agarose gel to visualise a product of ~600bp. 5ul of PCR1 product as template was amplified using unique i5 and i7 index primer combinations with 8 cycles and Q5 High-Fidelity DNA Polymerase (NEB) for each individual sample to allow pooling of sequencing libraries. The following primers were used:

FW:5’AATGATACGGCGACCACCGAGATCTACAC<u>**TATAGCCT**</u>ACACTCTTTCCCTACACGACGCTCTTCCGATCTtgtggaaaggacgaaaCACCG-3’

RV:5’CAAGCAGAAGACGGCATACGAGAT<u>**CGAGTAAT**</u>GTGACTGGAGTTCAGACGTGTGCTCTTCCGATCTATTTTA ACTTGCTATTTCTAGCTCTAAAAC-3’

The underlined bases indicate the Illumina (D501-510 and D701-712) barcode location that were used for multiplexing. PCR products were run on a 2% agarose gel, and a clean ~200bp band was isolated using Zymo Gel DNA Recovery Kit as per manufacturer instructions (Zymoresearch Inc.). Final samples were quantitated using Qubit dsDNA BR Assay and sent for Illumina Next-seq sequencing (20 million reads per 5 pooled lungs - 2×5 untreated and 2×5 treated) to the sequencing facility at Lunenfeld-Tanenbaum Research Institute (LTRI). Sequenced reads were aligned to sgRNA library using Bowtie version 1.2.2 with options –v 2 and –m 1. CRISPR screen hits were obtained and identified using <u>M</u>odel-based <u>A</u>nalysis of <u>G</u>enome-wide <u>C</u>RISPR-Cas9 <u>K</u>nockout (MAGeCK) algorithm.

### Analysis of genome editing efficiency

LSL-Cas9-GFP MEFs were cultured and infected with lentivirus carrying Cre and corresponding sgRNAs. Cells were live sorted for GFP expression and expanded further to extract genomic DNA using DNeasy Blood & Tissue Kit (Qiagen). Genomic DNA from tumors (GFP sorted and/or unsorted cells were used) from the mice injected with single sgRNAs was also isolated using the same kit. PCR was performed flanking the regions of sgRNA on genomic DNA from WT MEFs, cells infected with respective virus or tumors and sent for Sanger sequencing. Sequencing files along with chromatograms were uploaded to https://www.deskgen.com/landing/tide.html or https://ice.synthego.com/#/ and genome editing efficiency was estimated.

### *In vitro* T cell activation and expansion

Mouse T cells: Spleens were harvested from OT-I mice and dissociated to obtain single cell suspension. Red blood cells were lysed with ACK lysis buffer. Cells were resuspended at 1×10^6^ cells/ml in T cell media [RPMI-1640 (Gibco) + 10%FBS + 1% penicillin/streptomycin + 40μM 2-β-mercaptoethanol (Sigma-Aldrich)]. Medium was supplemented with 2μg/ml of SIINFEKL (OVA 257-264, AnaSpec) and human interleukin-2 (hIL-2, PeproTech) at 30 U/ml. After 2 days, equal amount of new medium supplemented with IL2 was added. Cells were used for in vitro assays following 4 days activation. For supplementary figure 3a: *Ex vivo* OT-I were isolated from spleen and LNs and purified using CD8a (Ly-2) microbeads (#130-117-044, Miltenyi Biotech). CD8^+^ OT-I cells were labeled with 1μM CFSE (CellTrace™ CFSE proliferation kit, Molecular Probes) and cultured in triplicates with plate-bound anti-CD3 (5 μg/ml) and anti-CD28 (2 μg/ml) mAb in 200ul of T cell media supplemented with 5 ng/ml of IL-7 and 30 U/ml of IL-2 (PeproTech) for 4 and 7 days. Activation and proliferation of CD8^+^ OT-I cells were determined by flow cytometry.

Human T cells: As described previously, activated and expanded day 15 human DNT (γδ) cells were kindly provided by Dr. Li Zhang’s lab (Toronto General Hospital, UHN)^103^. Briefly, blood samples were obtained from healthy donors upon consent with a protocol approved by the UHN Research Ethics Board. DNTs were enriched by depleting CD4^+^ and CD8^+^ cells using RosetteSep™ human CD4- and CD8-depletion cocktails (Stemcell Technologies). The CD4 and CD8 depleted cells were cultured in 24-well plates pre-coated with 5◻μg/ml anti-CD3 antibody (OKT3, eBioscience) for 3◻days in RPMI-1640 (Gibco) supplemented with 10% FBS (Sigma) and 250◻IU/ml IL-2 (Proleukin). Fresh IL-2 and OKT3 were added to the DNT cultures every 2–4◻days. After 10 days of activation/expansion, 0.1ug/ml of OKT3 250◻IU/ml IL-2 was added to culture every 2-4 days. DNTs were harvested between day 15–20 and purity was assessed by flow cytometry prior to experiments. The mean purity of DNTs used in the study was ~94%, and cells were used for functional studies.

### Synthetic Peptides and inhibitors

Synthetic peptides were generated by AnaSpec and purified with HPLC to ≥ 95 % purity and verified by Mass Spectrometry. H2K^b^-restricted peptide epitope of OVA (257-264) - SIINFEKL were dissolved in PBS at a concentration of 2 mg/ml. For *in vivo* experiments mice were i.p. immunized with 100μg of OVA (SIINFEKL) emulsified in Complete Freund’s adjuvant (CFA) or Incomplete Freund’s adjuvant (IFA). Blocking antibodies CTLA-4 (clone 9H10), PD-L1 (clone 10F.9G2) were obtained from BioXcell and administered every other day throughout the experiment at a dose of 150 μg per mouse per treatment. All procedure were performed in accordance with TCP SOP #SAF034.

### Adoptive T cell transfer (ACT)

For supplementary figure 3b-d: CD8 T cells were isolated from gender-matched OT-I spleen and LNs using CD8a (Ly-2) microbeads (#130-117-044, Miltenyi Biotech). 10×10^6^ OT-I cells were IV injected (group 2 and 3) into C57BL6/J mice followed by i.p. injection of 100μg of OVA (SIINFEKL) emulsified in CFA and IFA (d7). Spleens were isolated 3 days after immunization. Activation and proliferation of CD8^+^ OT-I cells were determined by flow cytometry.

For lung cancer treatments: ACT of gander-matched, purified 15-20×10^6^ OT-I cells were i.v. transferred into 3-4 weeks old LSL-Kras ^G12D^-CAS9 -LUC or LSL-Braf ^V600E^ -CAS9-LUC mice after lung tumor induction, which was assessed by bioluminescence imaging (BLI). Mice were immunized on day 1 (100μg of OVA in CFA) and day 7 (100μg of OVA in IFA) after ACT. In some experiments, spleen, LNs, and/or lungs were isolated for total cell number enumeration and flow cytometric analysis on day 7 or at endpoints. For tumour tissues, the entirety of each sample was acquired and the total number of CD3^+^CD8^+^Tom^−^ T cells and transferred OT-I cells was assessed. In some experiments lungs were perfused with 4% PFA and isolated for H&E staining and immunofluorescence analysis. All procedure were performed in accordance with TCP SOP #SAF034.

### *In vitro* killing assay

#### Mouse T cells

4 days activated and expanded splenocytes (effectors, E) from OT-I crossed to mT/mG mice were co-cultured with pre-plated 20,000 LLC transduced with either sgNTC, sg sg*Serpinb9,* LLC-CTLR-SIINFEKL or ADAM2-SIINFEKL cells (targets, T) and labeled with 1μM CFSE (Molecular Probes) in triplicates in 48-well plates at varying E:T ratios for 4 hours. To determine the cytotoxicity induced by CD8 T cells, drop in CFSE (as read out of targets being killed) was measured by flow cytometry. Following formula was used: percent of killing = 100 - total number of PI^−^CFSE^+^ cells with effectors/total number of PI^−^CFSE^+^ without effectors x 100%.

#### Human T cells

Donor-derived (UPN119, UPN133) day 15-20 activated and expanded DNT (yO) T cells (effectors, E) were co-cultured with CFSE (1μM) labeled 20,000 pre-plated human lung cancer cells (A549-*SERPINB9* or H125-*SERPINB9*) in triplicates in 48-well plates at varying E:T ratios for ~16-18 hours (overnight). The cytotoxicity induced by DNTs was determined by flow cytometry where percent of killing = 100 -total number of PI^−^CFSE^+^ cells with DNTs/total number of PI^−^CFSE^+^ without DNTs x 100%.

### Western blotting

Cell lysates were generated using RIPA lysis buffer supplemented with protein inhibitor (Roche, #4693159001) and protein concentration was determined using Pierce™ BCA protein assay. Denaturated lysates (20ug) were applied to a 4-15% Mini-PROTEAN TGX precast protein gels (Biorad) or to a 10% SDS-PAGE and blotted using standard procedures. For protein detection, blots were incubated with primary V5 tag monoclonal (#R960-25, Life Technologies) antibody overnight and with secondary antibody (goat anti-mouse IgG-HRP (#1706516, Biorad) for 1h. Direct-Blot™ HRP anti-GAPDH (#607904, Biolegend) was used as a loading control. Chemiluminescence was used to visualize the protein bands (Biorad). In some experiments tumors were first sonicated in RIPA buffer and protocol described above for western blotting was followed.

### Tumor implantation

Mice were anesthetised with 2-2.5% isoflurane with oxygen. LLC-CTLR-SIINFEKL or LLC-ADAM2-SIINFEKL cell lines were gently injected subcutaneously in the upper right backside of mice. 1×10^5^ cells were injected per mouse for all the experiments. After tumor onset, tumor growth was measured every day or every second day by digital caliper. Intravenous injection of 15-20×10^6^ OT-I was performed on D19 after tumor reached the volume between 100-600 mm^3^, followed by immunization with OVA/CFA on day 1 and OVA/IFA on day 7. Humane intervention points were called when tumor size reached 1700 mm^3^, according to SOP AH009.

### Preparation of tumor tissues for flow cytometry

Tumor tissues (fresh or frozen) were minced into small pieces using surgical blade and scalpel (08-957-5D, Fisher Scientific) and processed using the tumor dissociation kit (130-096-730, Miltenyi Biotec) as recommended by supplier. Single-cell suspensions from lung or implanted tumors were obtained using kit and the gentleMACS Octo Dissociator. Dissociated cells were passaged through 70μm cell strainer (BD), collected in a 50ml falcon tube and resuspended in staining buffer (1%BSA in PBS).

### Flow cytometry

Single cell suspensions from spleen, LNs, lung or implanted tumors were washed with FACS buffer (DPBS + 1%BSA+ 1mM EDTA + 0.1% sodium azide), incubated with FC block (CD16/32) for 30’ at 4°C, stained with appropriate antibodies and washed twice in FACS buffer. Dead cells were excluded from all data by forward and side scatter and 4’,6-diamidino-2-phenylindole, dihydrochloride (DAPI, Molecular Probes - 5mg/ml, used 1/50,000) or fixable viability dye eFluor™ 450 (1/1000, Invitrogen). Cells were stained by standard staining techniques and analyzed on Fortessa flow cytometer (BD Biosciences). For intracellular staining, the cells were permeabilized using eBioscience™ Foxp3/Transcription factor Staining Buffer Set (00-5523-00,Thermo Fisher Scientific) according to manufacturer’s recommendation. Data Files were analyzed using Flow-Jo (Tree Star).

### Immunofluorescence

Cells or tissue sections were fixed with 4% paraformaldehyde for 10 minutes. Following fixation, slides were rinsed 3 times with PBS for 5 minutes, permeabilized using 0.5% Tween-20 in PBS at 4ºC for 20-minutes and rinsed with 0.05% Tween-20 in PBS for 3×5 minutes at room temperature. Samples were blocked at room temperature with blocking serum (1% BSA, 1% gelatin, 0.25% goat serum 0.25% donkey serum, 0.3% Triton-X 100 in PBS) for 1 hour. Samples were incubated with primary antibody diluted in blocking serum overnight at 4ºC followed by 3 washes for 5 minutes in PBS. Secondary antibody was diluted in blocking serum with DAPI and incubated for 1 hour at room temperature in the dark. Following incubation, samples were washed 3 times for 5 minutes in PBS. Coverslips were added on slides using MOWIOL/DABCO based mounting medium and imaged under microscope next day. For quantification, laser power and gain for each channel and antibody combination were set using secondary antibody only as control and confirmation with primary positive control and applied to all images. Images were captured and expression of the specified genes were quantified with fluorescent Nikon eclipse Ti inverted microscope.

### RNAscope

Custom-designed 20 ZZ probe targeting 1279-2224 bp of NM_009618.3 mouse Adam2, named mm-Adam2; catalogue probes eGFP (cat # 538851-C1) and TOM (cat # 317041-C2) were ordered from Advanced Cell Diagnostics, ACD. Adam2, eGFP, Tom in situ hybridization was measured by RNAscope assay (Advanced Cell Diagnostics, ACD) according to the manufacturer’s protocol using RNA scope^R^ Multiplex Fluorescent Reagent kit v2 (323-100). Briefly, paraffin embedded normal (negative ctlr) and tumor-bearing lungs isolated from LSL-Kras ^G12D^-CAS9-LUC mice were cut into 5μm sections and hybridized at 40°C for 2 hours. Adam2 ORF expressing tumor bearing lungs or normal testis were used as positive controls. Hybridization signal was amplified using AMP 1, 2, 3 and developed using appropriate HRP signal. Images were captured with fluorescent Nikon eclipse Ti2 inverted microscope.

### RNA sequencing

Total RNA was prepared from tumor tissues using TRIzol reagent (Invitrogen) or the Quick-RNA MiniPrep kit (R1055, Zymo Research) treated with ezDNase (Invitrogen). The RNA samples were quality checked by LTRI Sequencing facility using 5200 fragment analyzer system, with all samples passing the quality threshold of RQN score >9.3 except two samples with a score of 7.6. The library was prepared using an Illumina TrueSeq mRNA sample preparation kit at the LTRI sequencing facility, and complementary DNA was sequenced on an Illumina Nextseq platform. Sequencing reads were aligned to mouse genome (mm10) using Hisat2/bowtie2 version 2.1.0 and counts were obtained using featureCounts (Subread package version 1.6.3). Differential expression was performed using DEseq2 release 3.8. Gene set enrichment analysis was performed using GSEA computational method software released by Broad institute.

