## Supplementary Figures for "*In vivo* CRISPR screens reveal Serpinb9 and Adam2 as regulators of immune therapy response in lung cancer"

A.

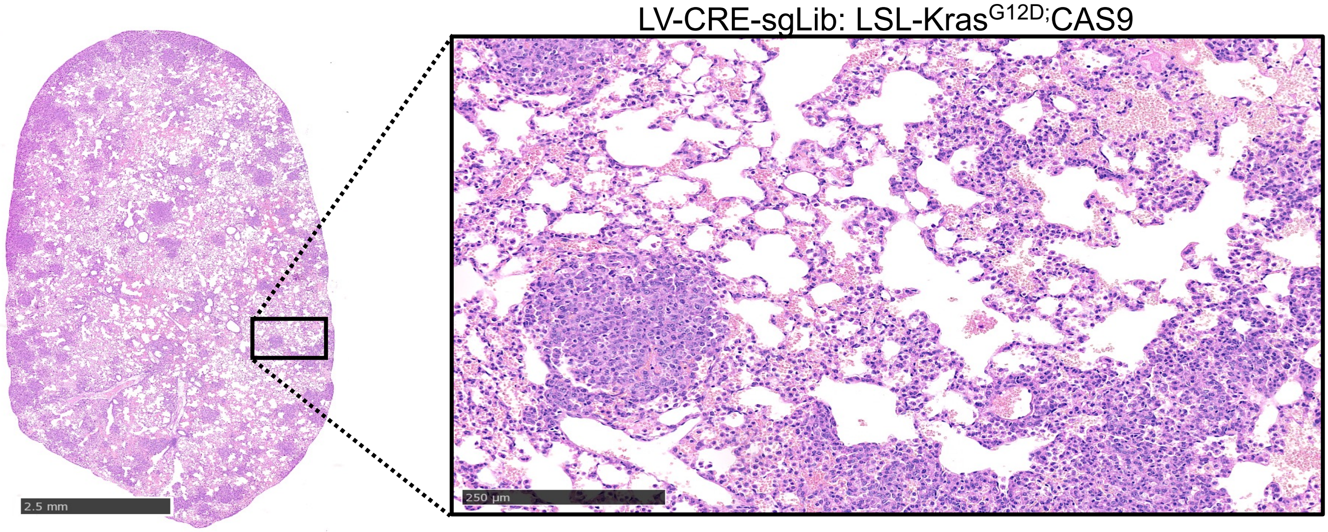


B.

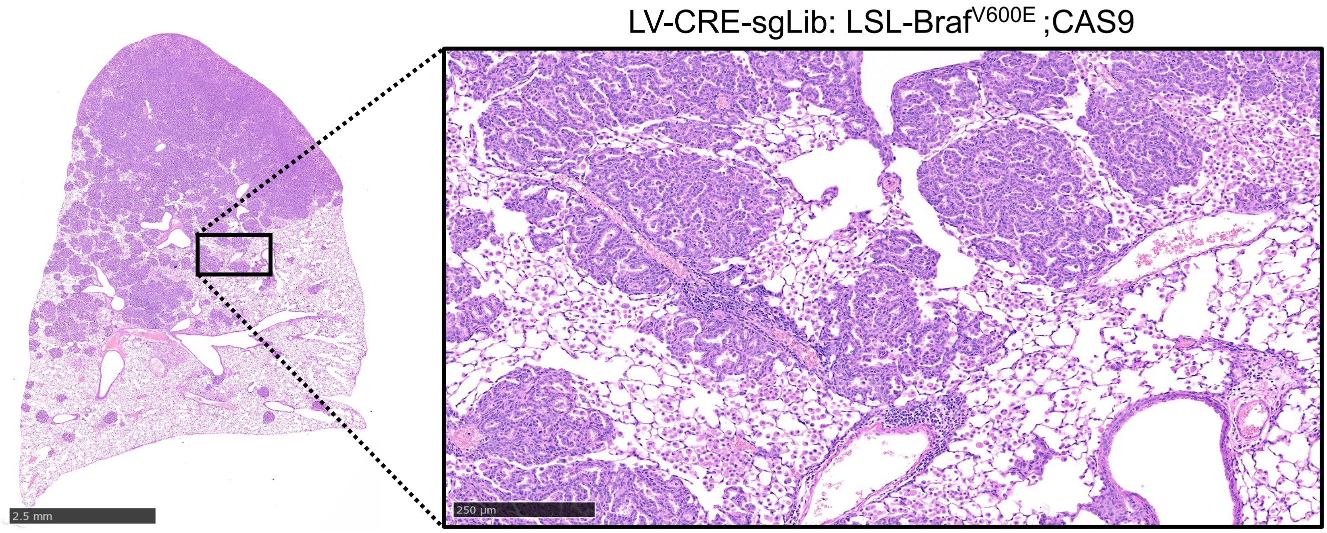


C.

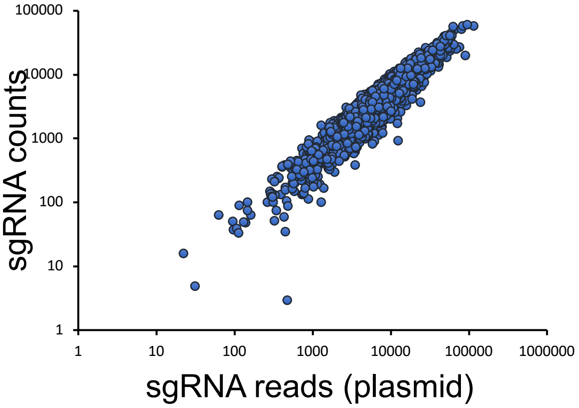


**Supplementary Figure 1**

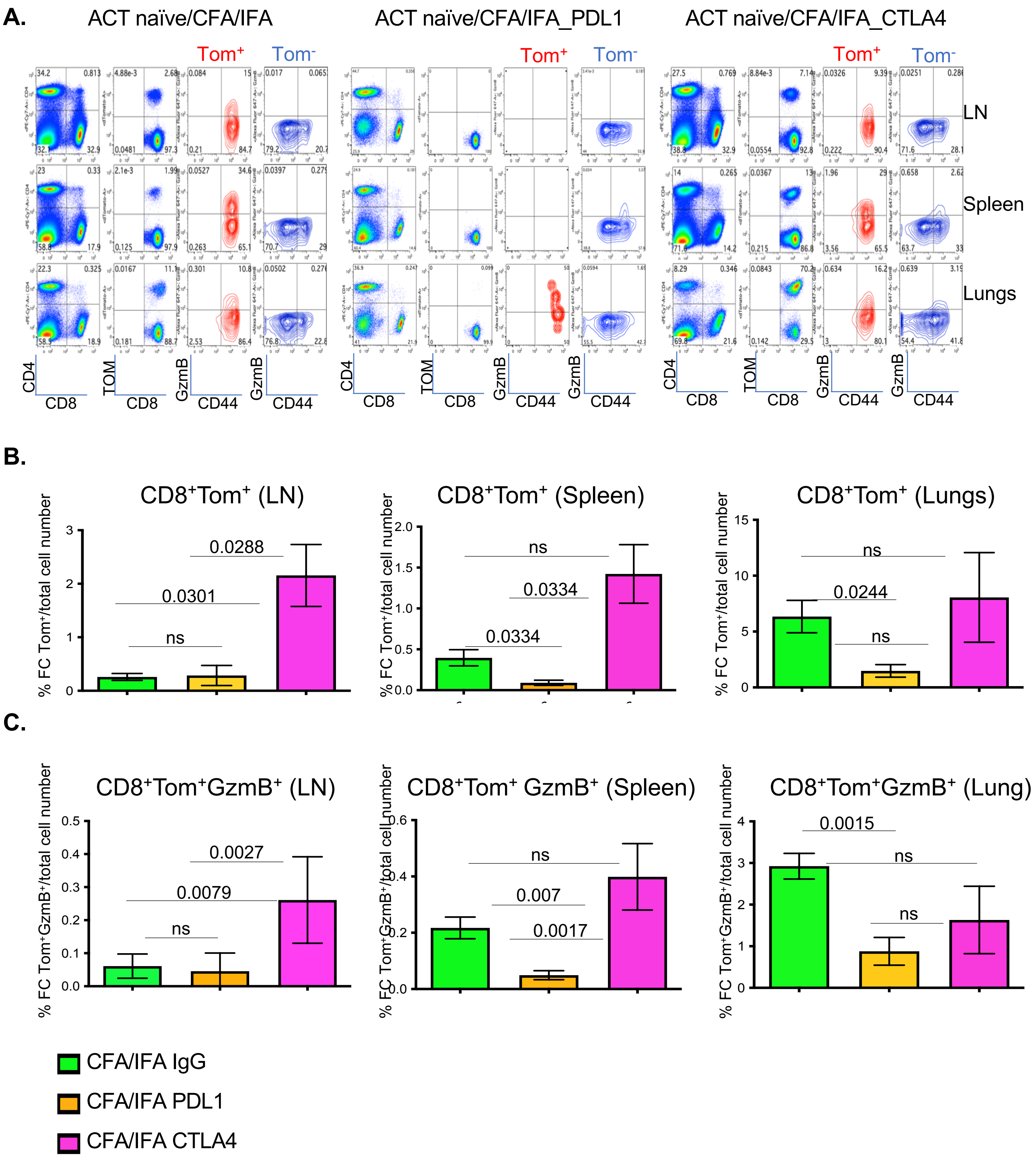


**Supplementary Figure 2**

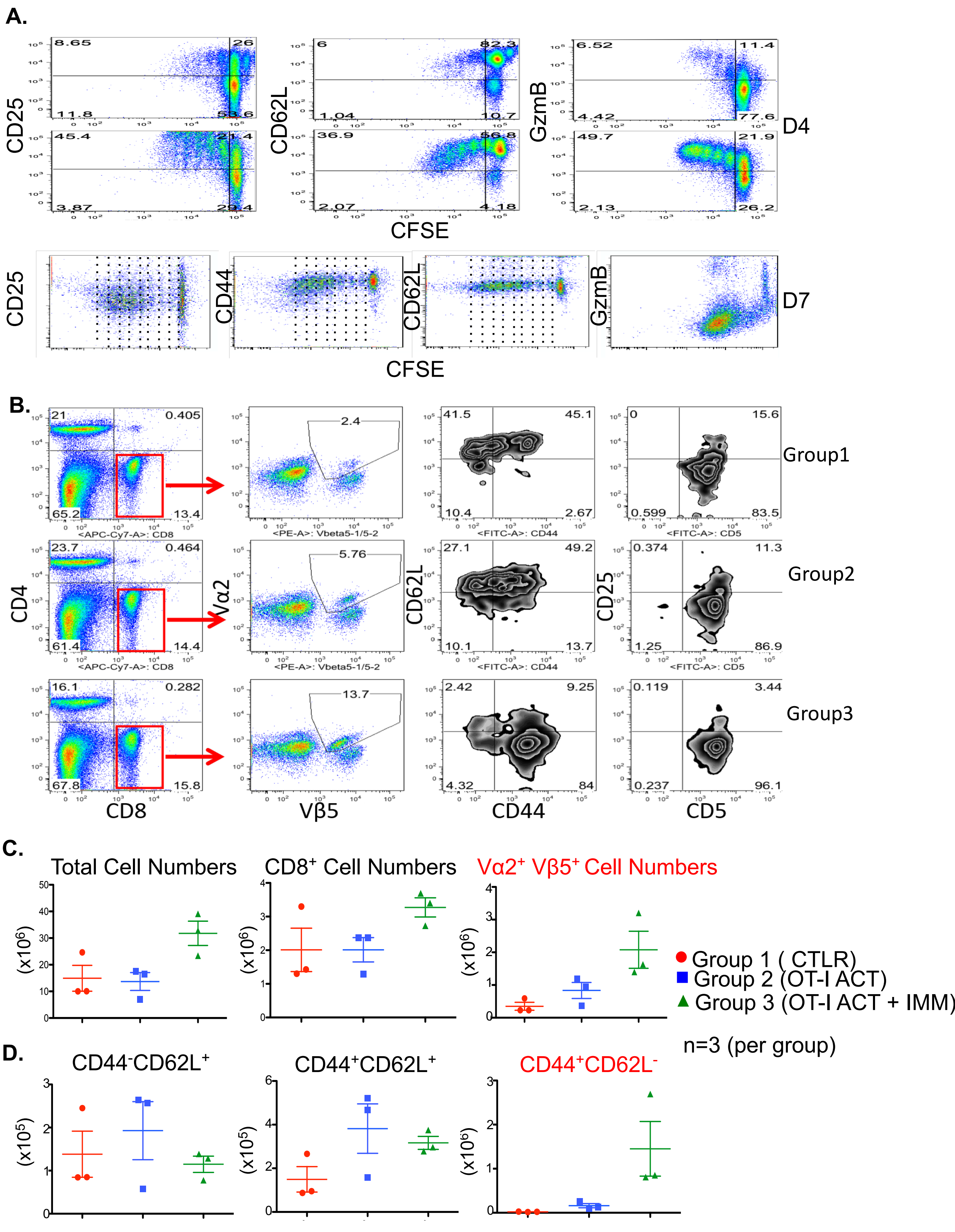


**Supplementary Figure 3**

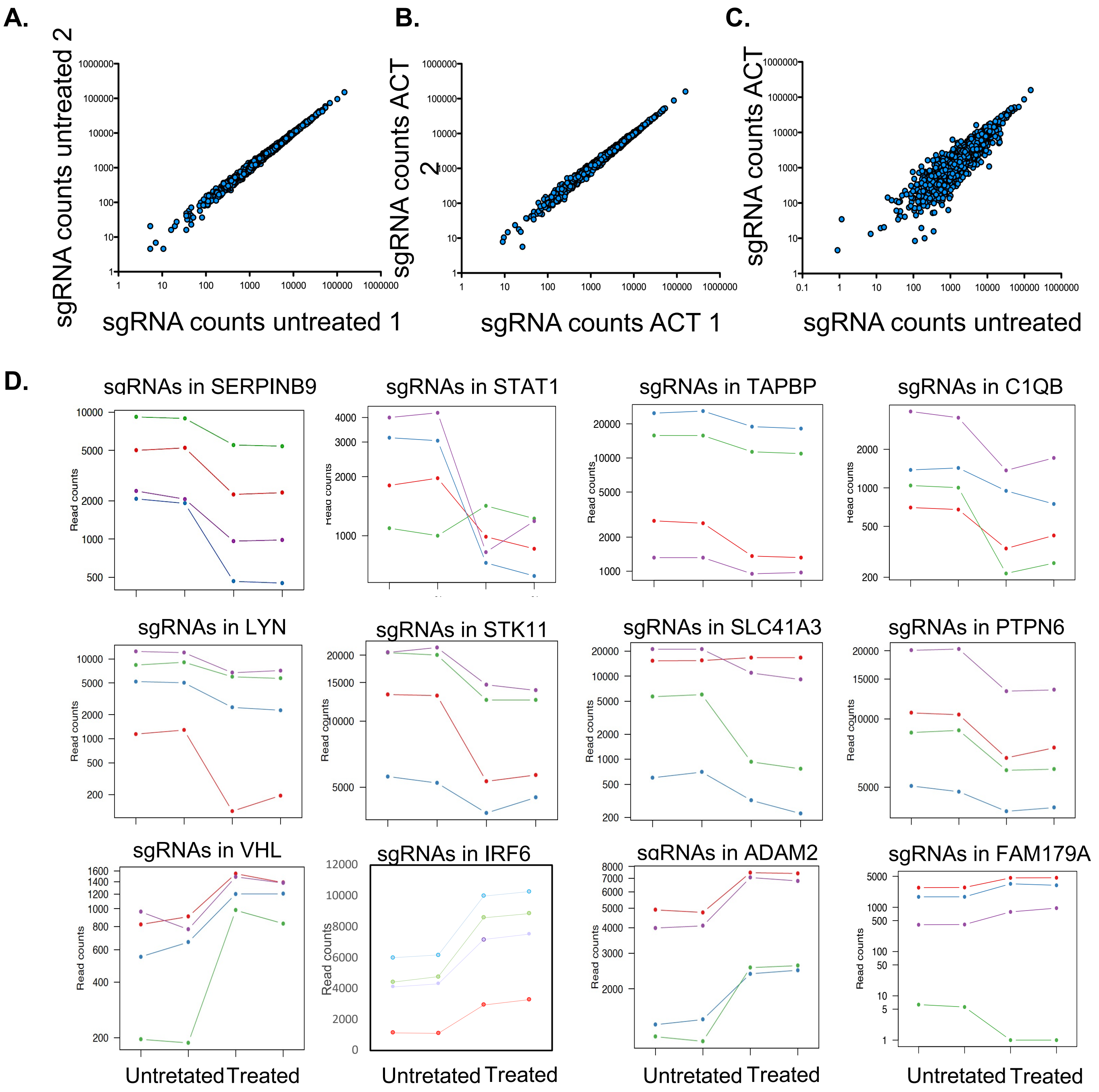


**Supplementary Figure 4**

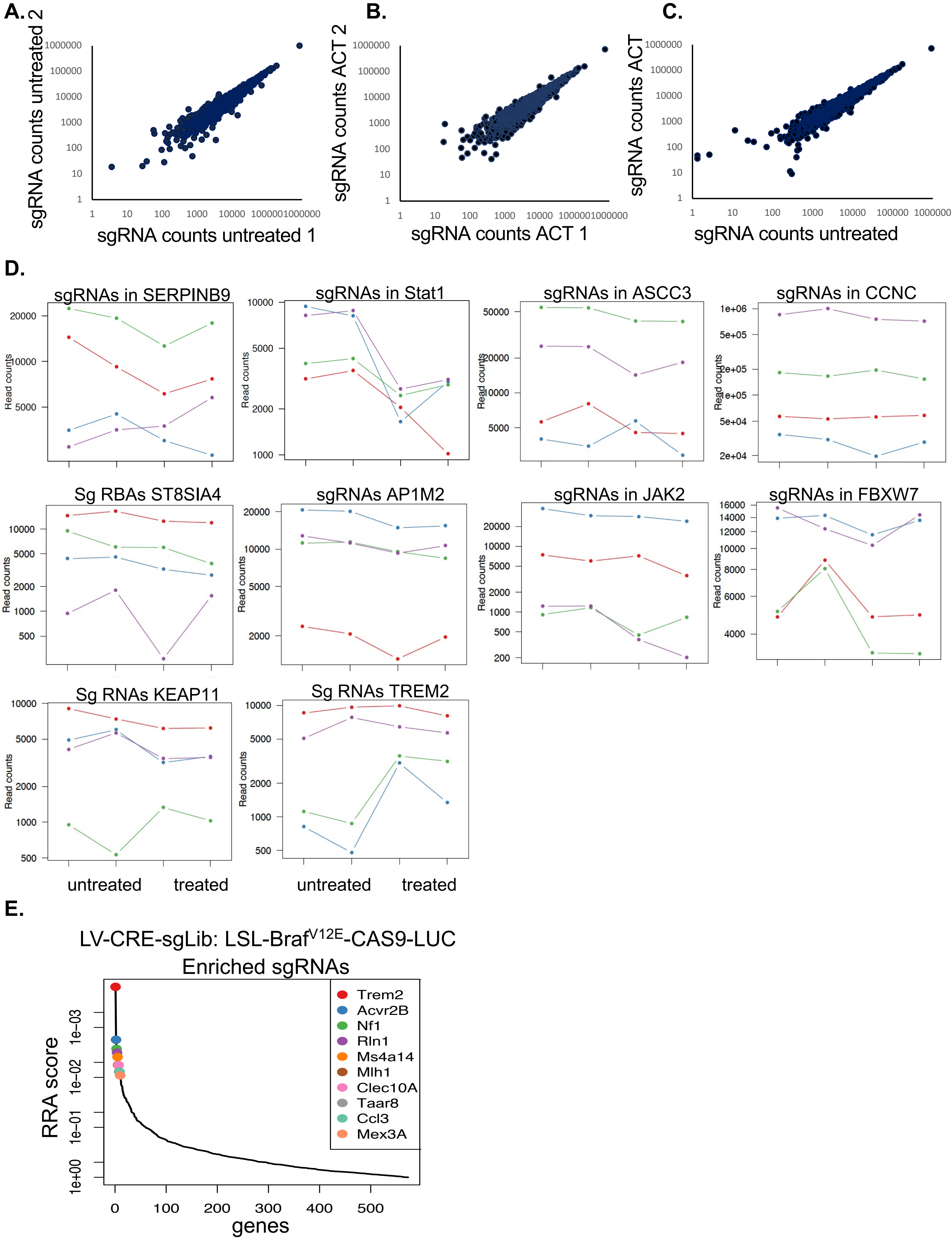


**Supplementary Figure 5**

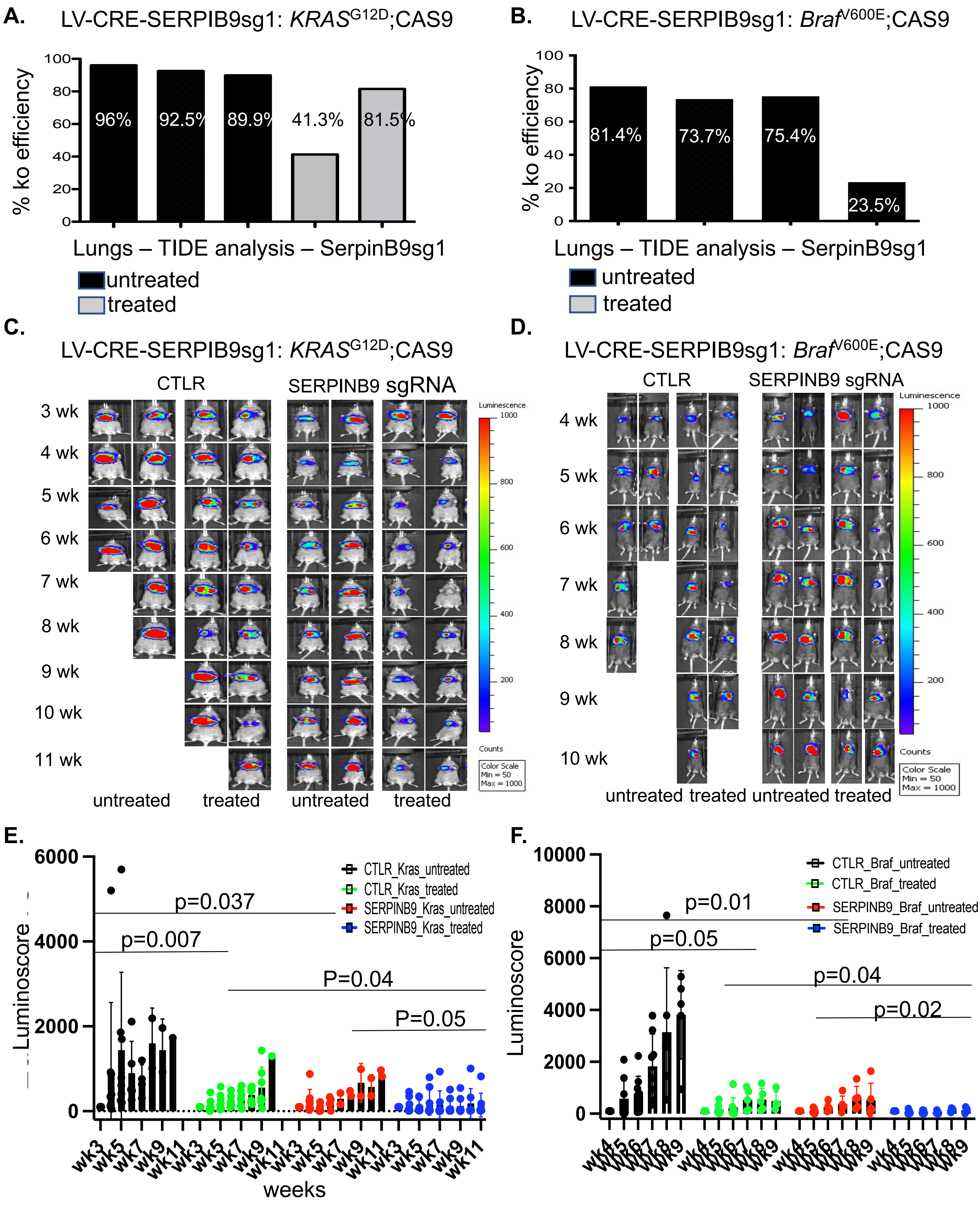


**Supplementary Figure 6**

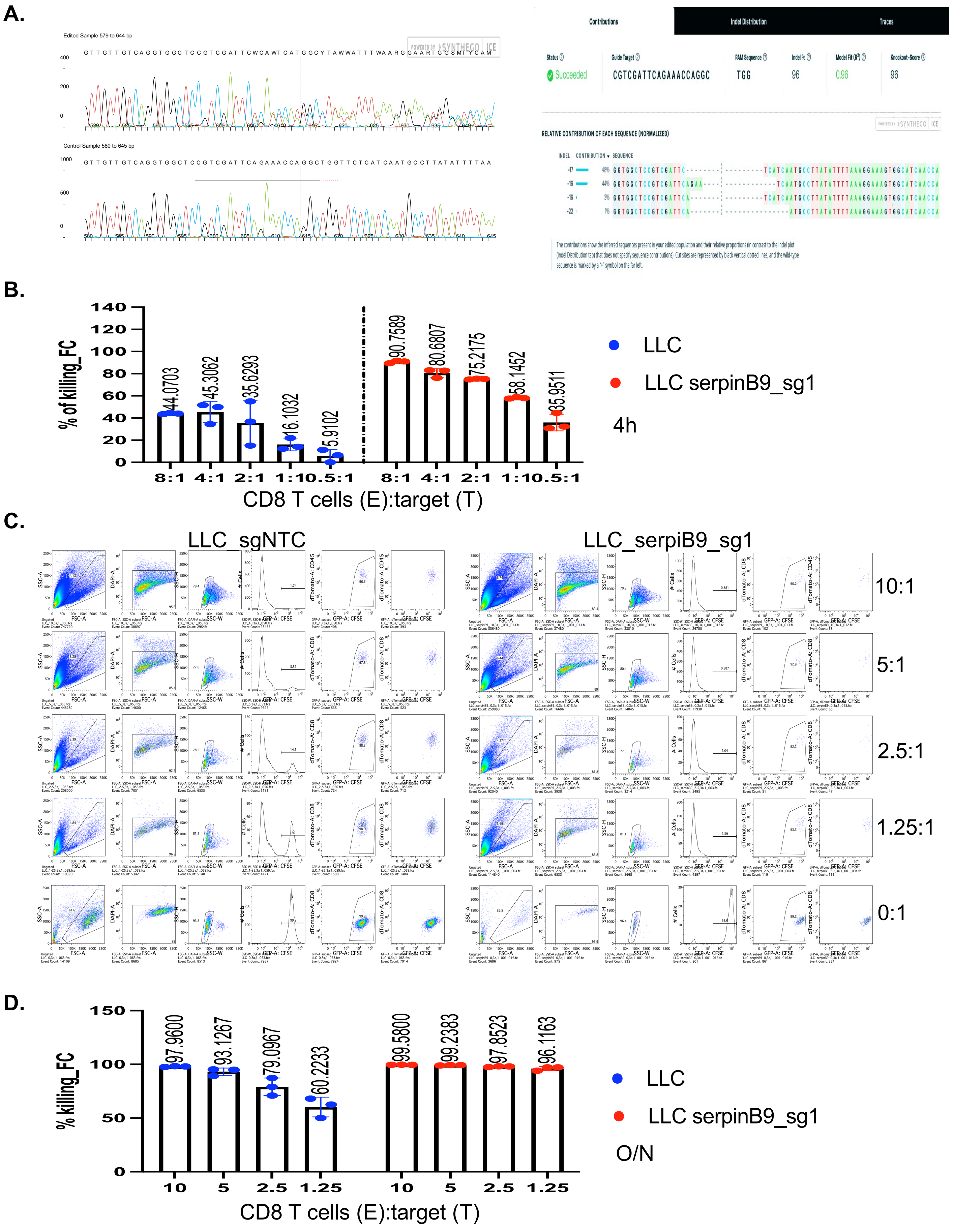


**Supplementary Figure 7**

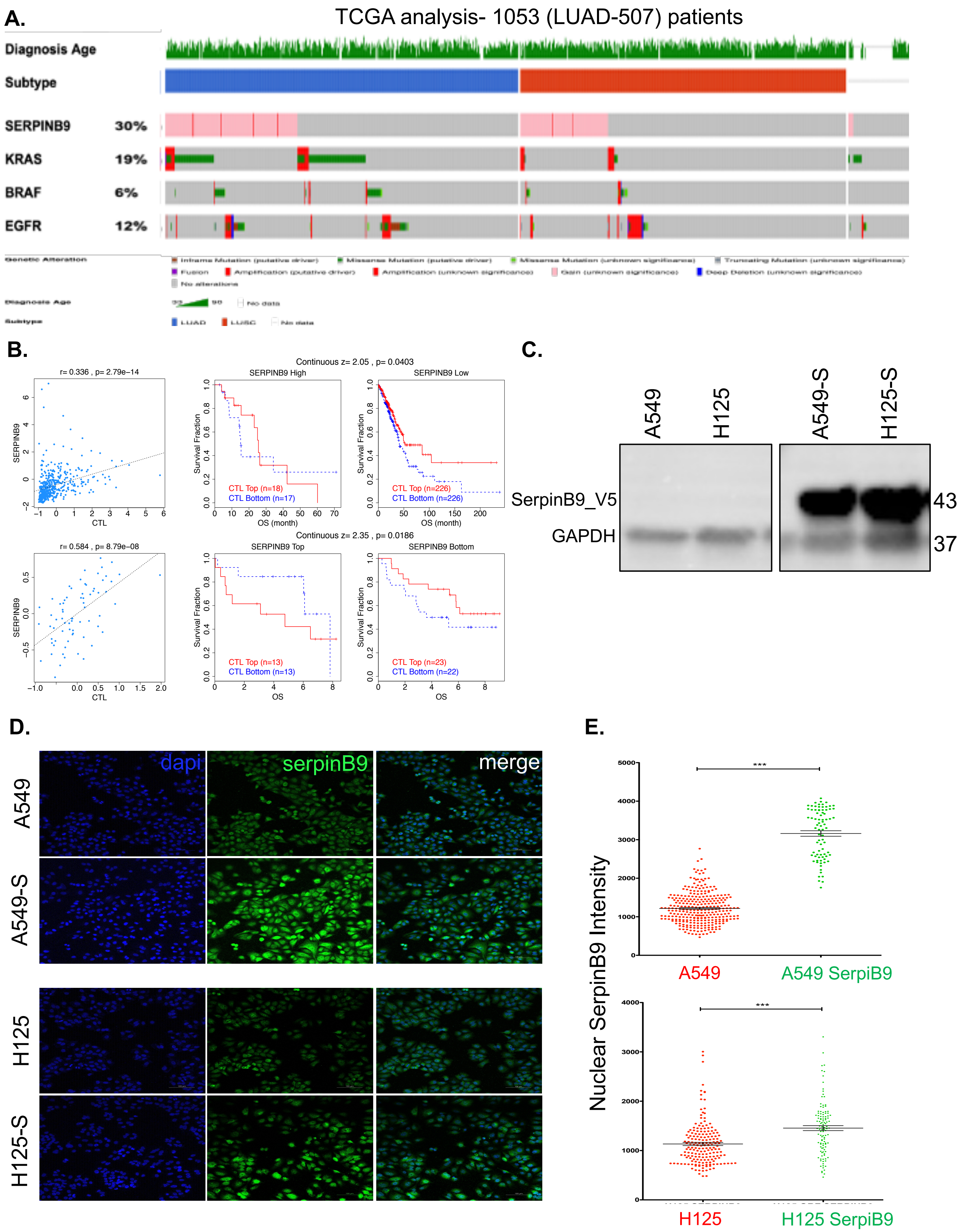


**Supplementary Figure 8**

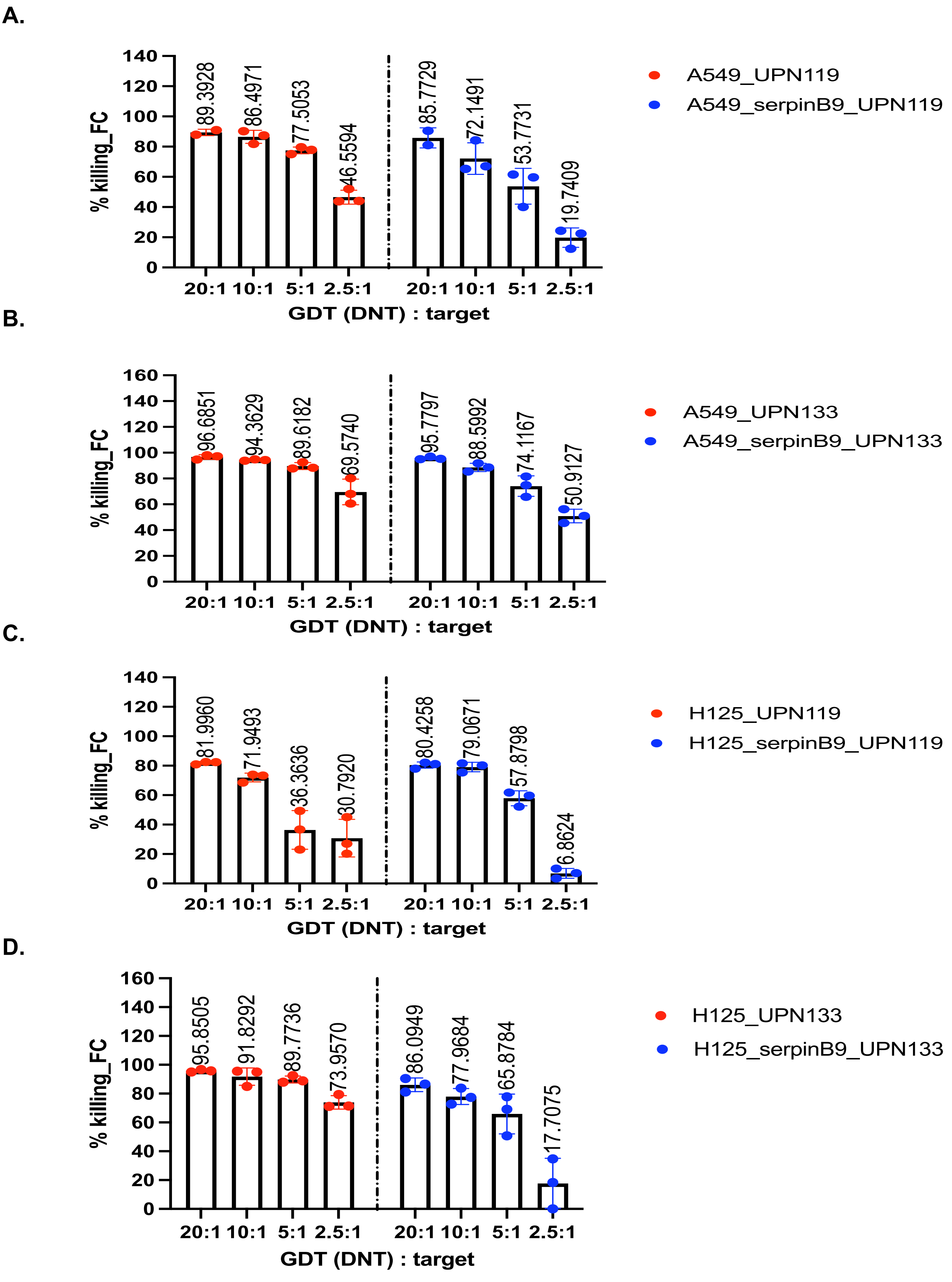


Supplementary Figure 9

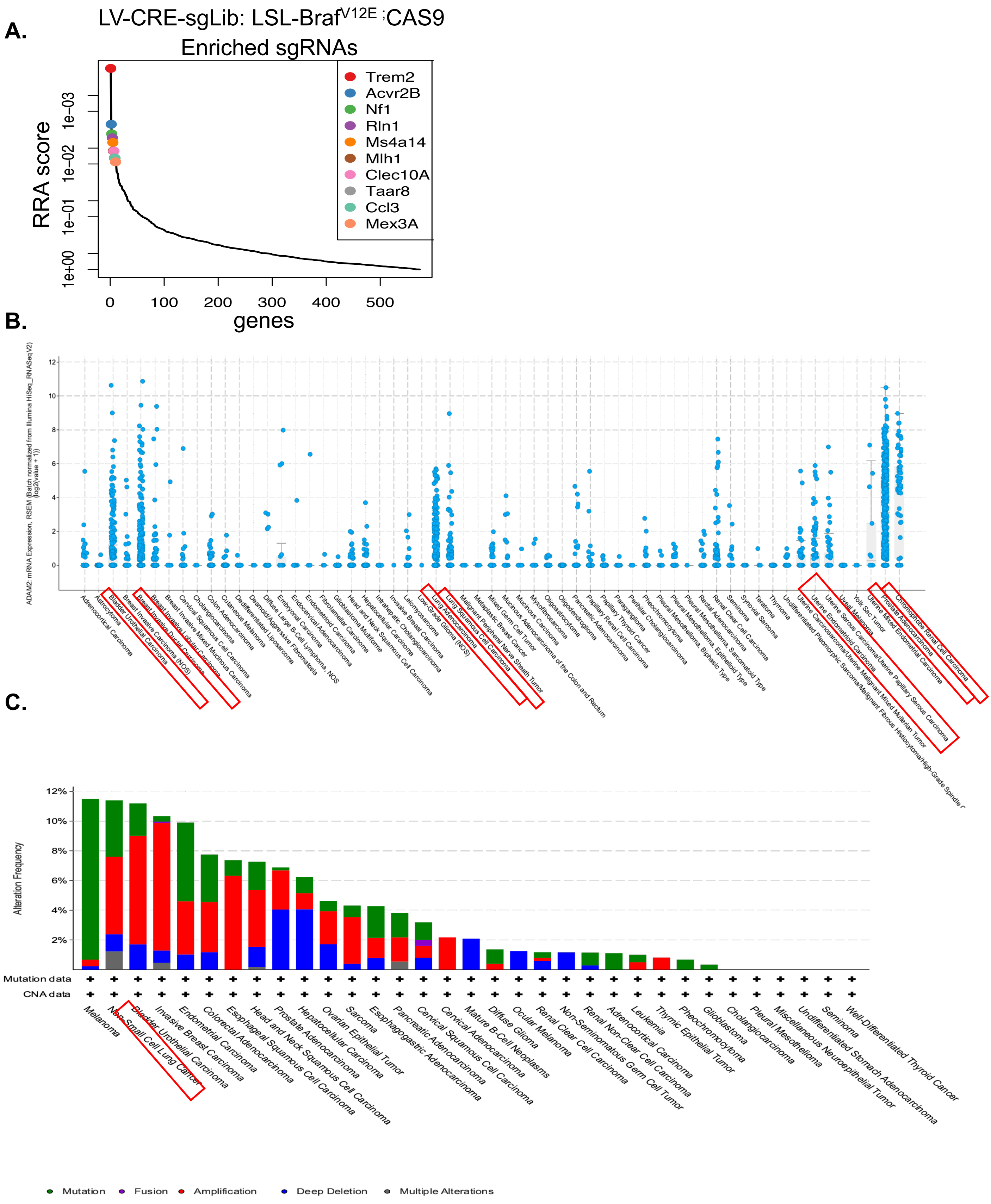


**Supplementary Figure 10**

A.

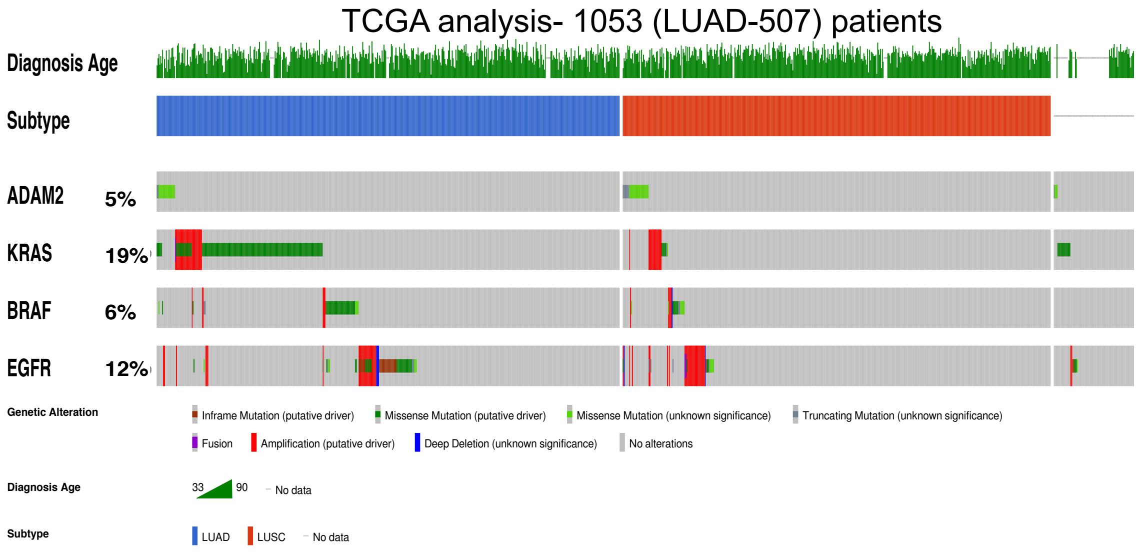


B.

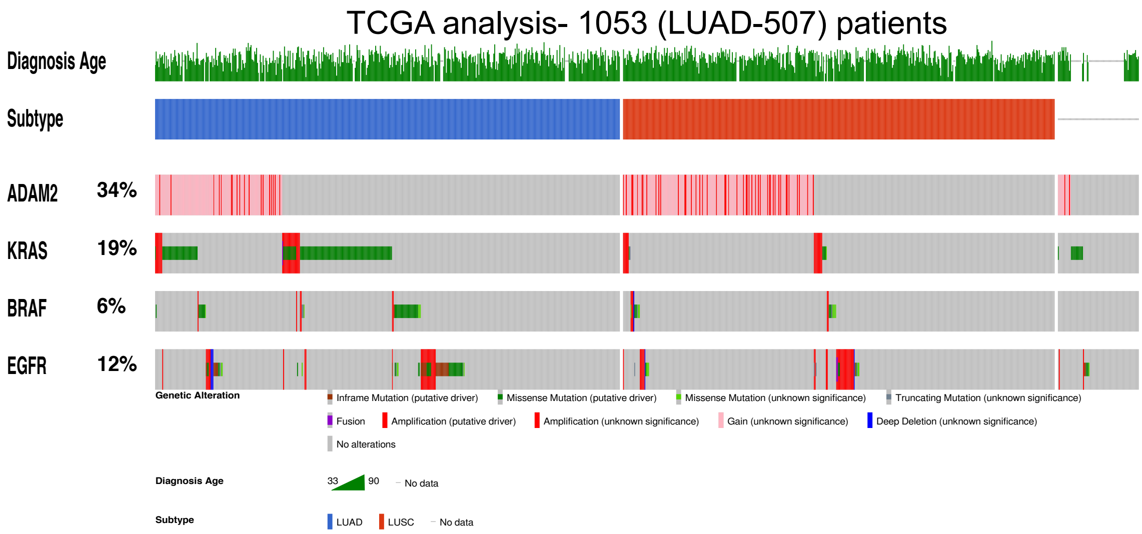


C.

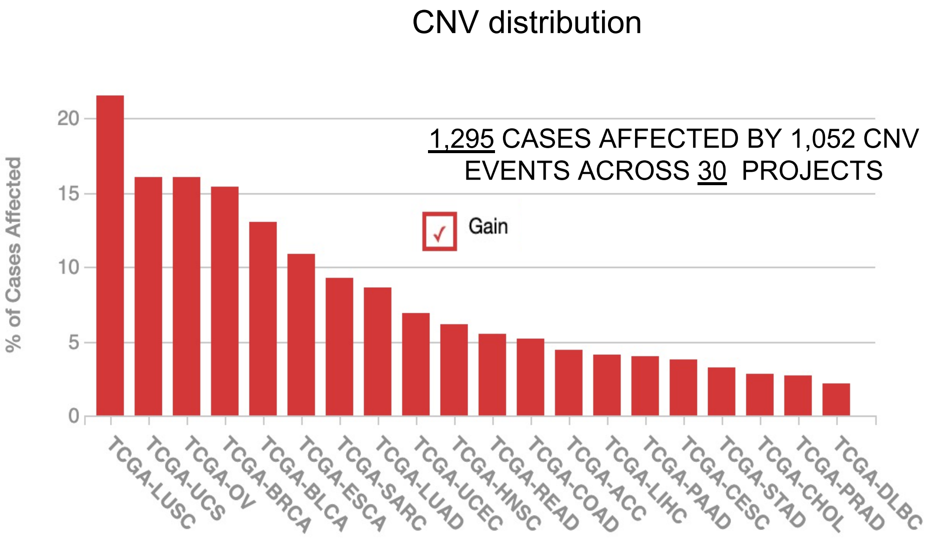


**Supplementary Figure 11**

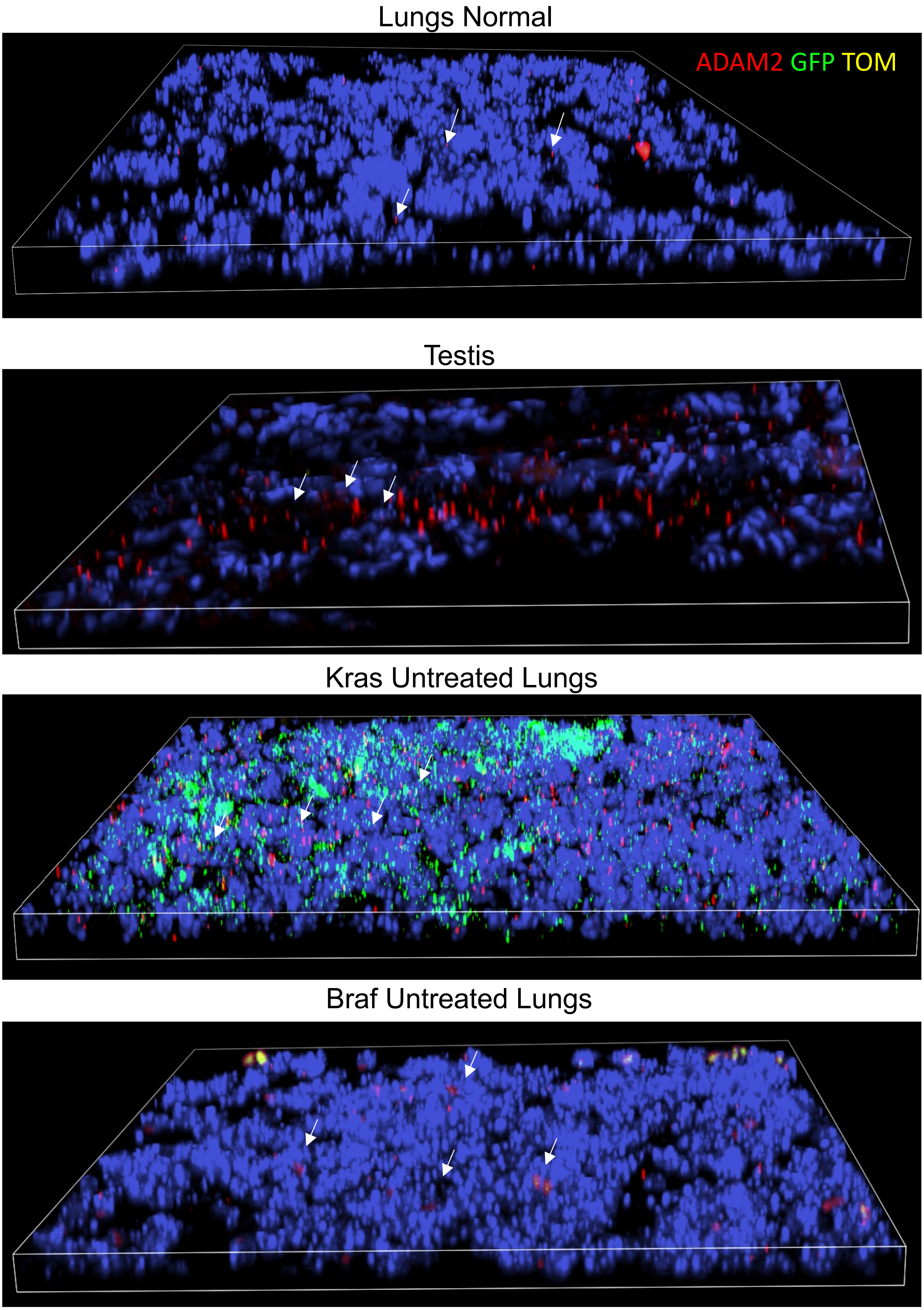


**Supplementary Figure 12**

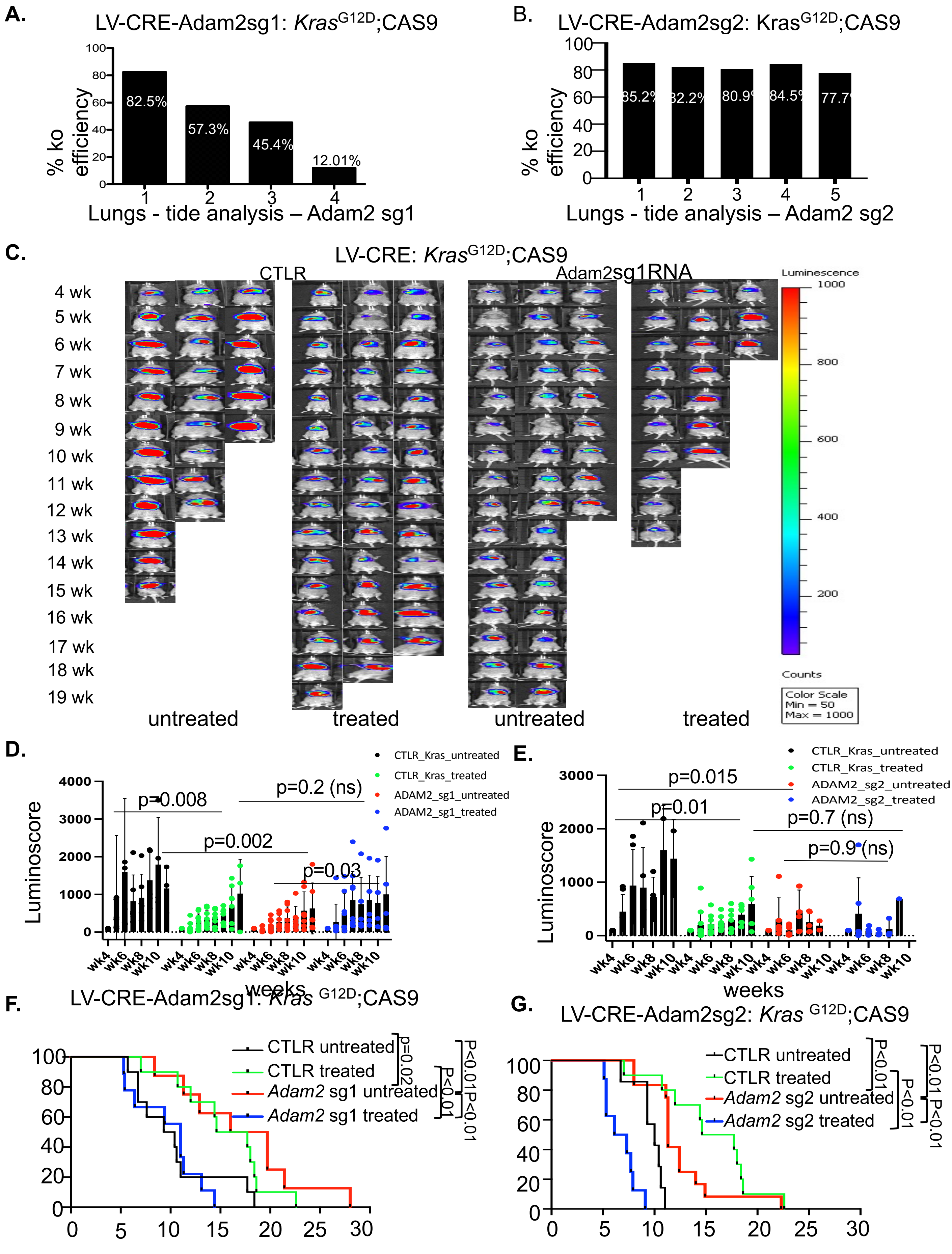


**Supplementary Figure 13**

**A.**

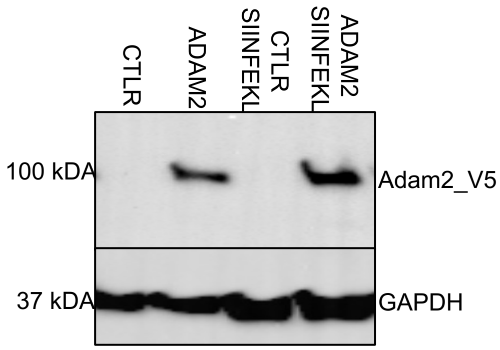


**B.**

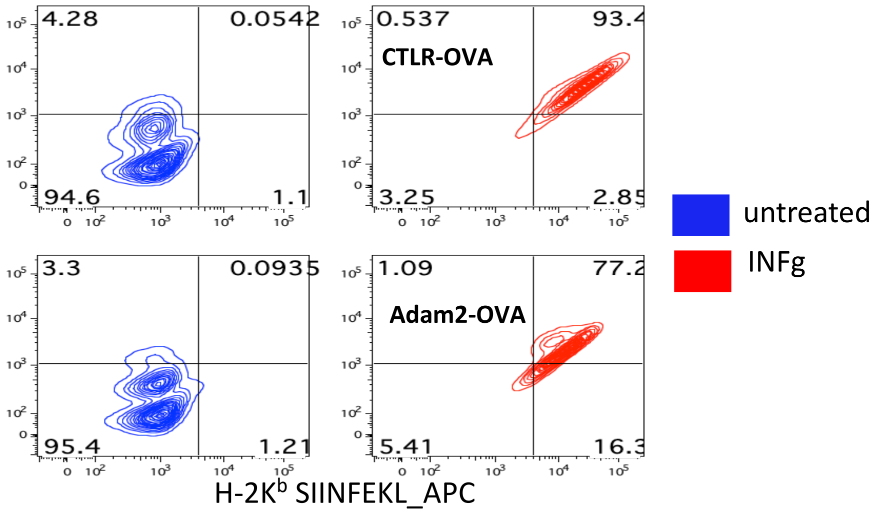


**Supplementary Figure 14**

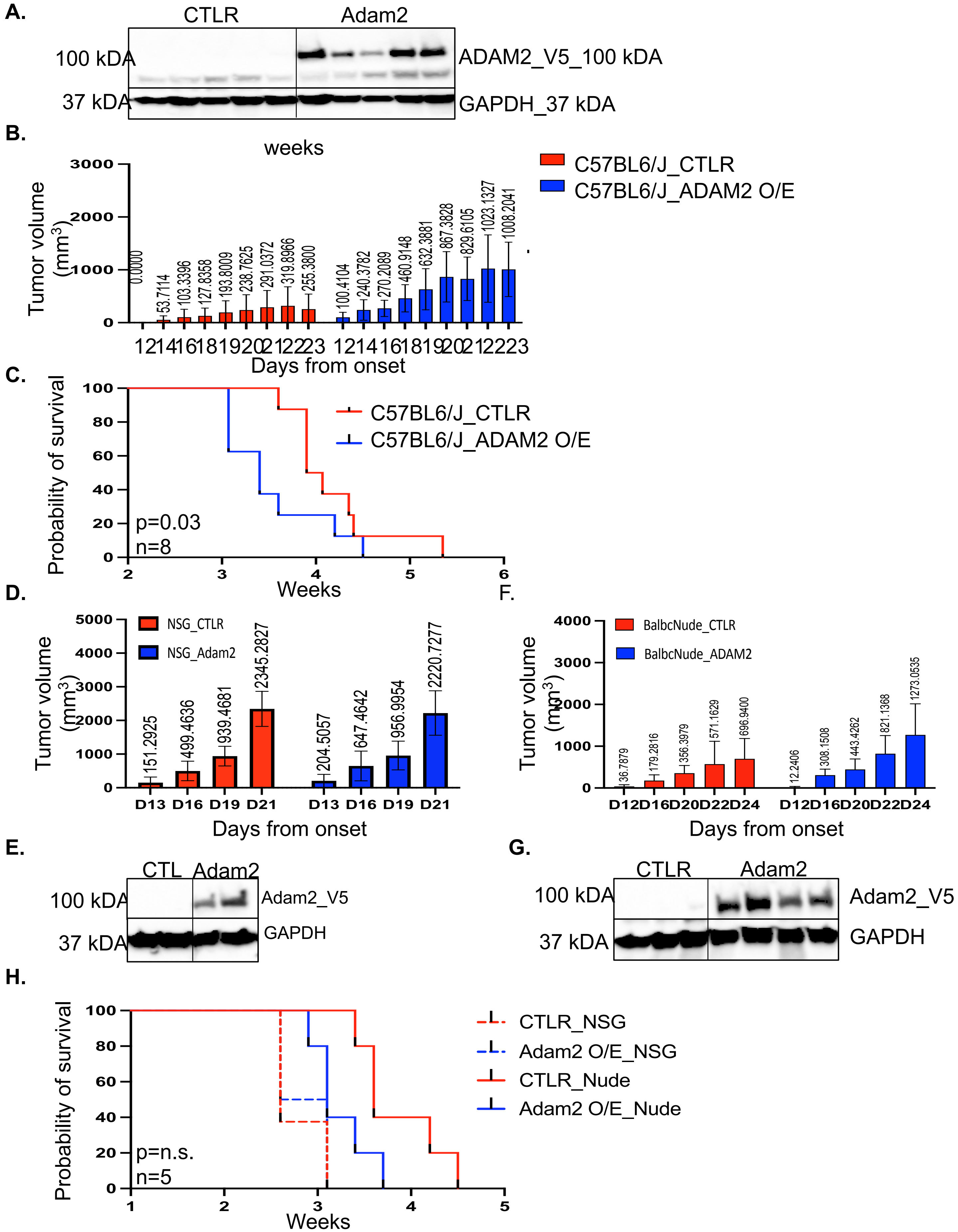


**Supplementary Figure 15**

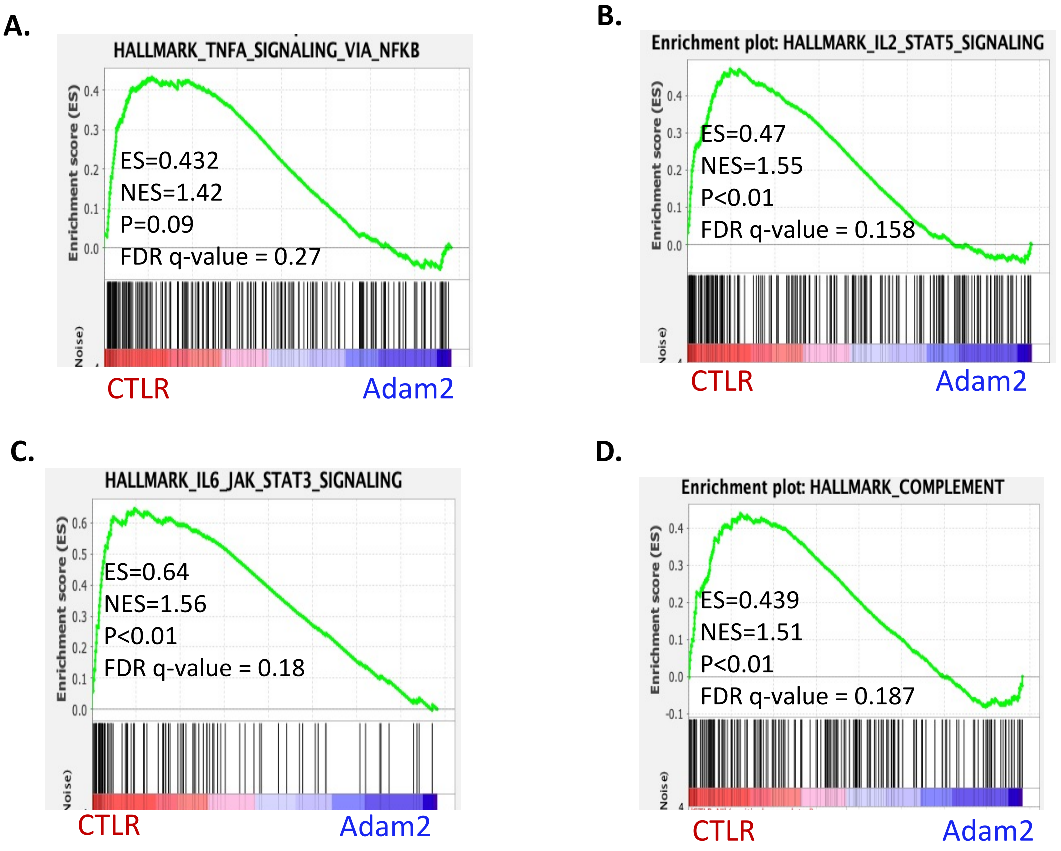


Supplementary **Figure** 16

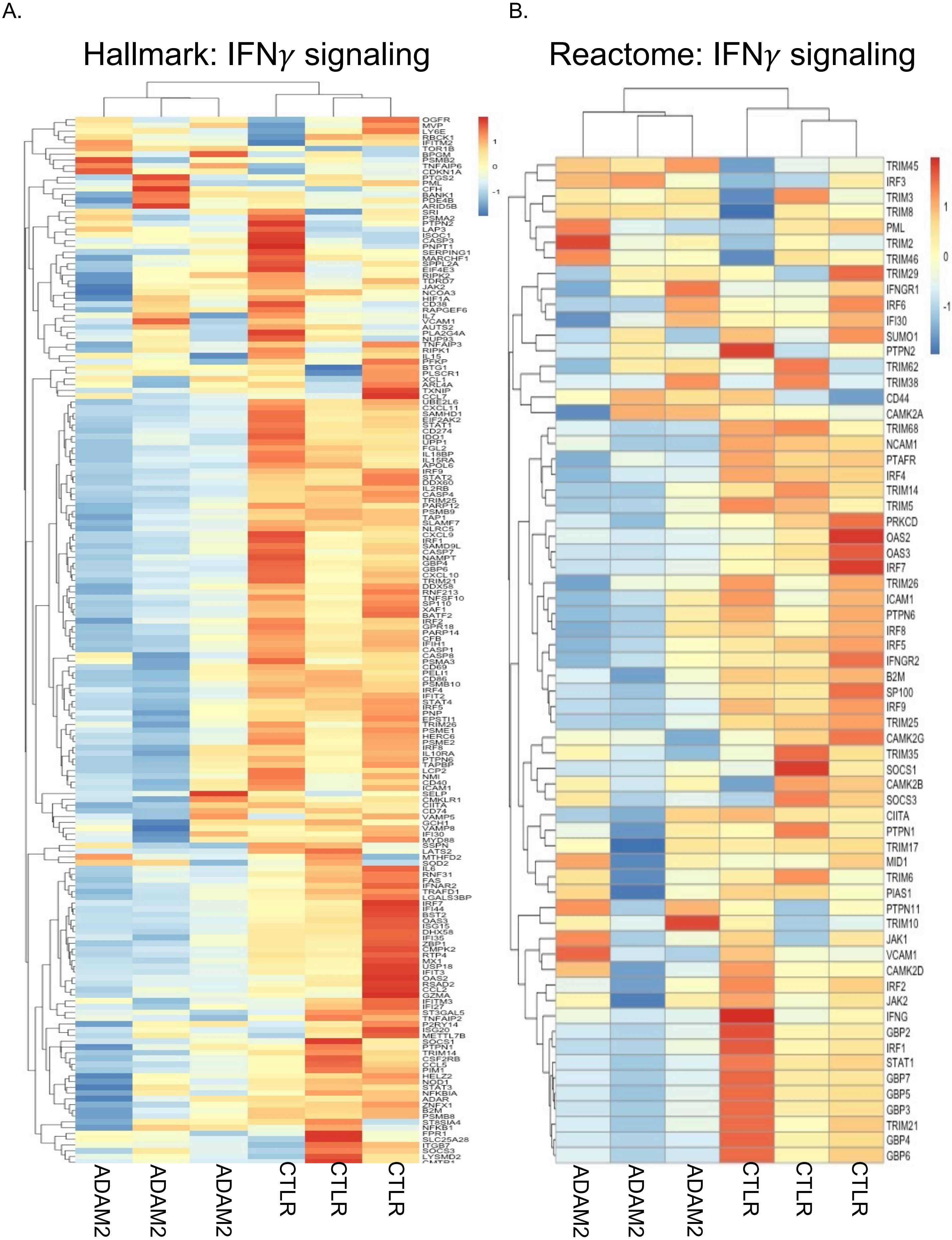


**Supplementary Figure** 17

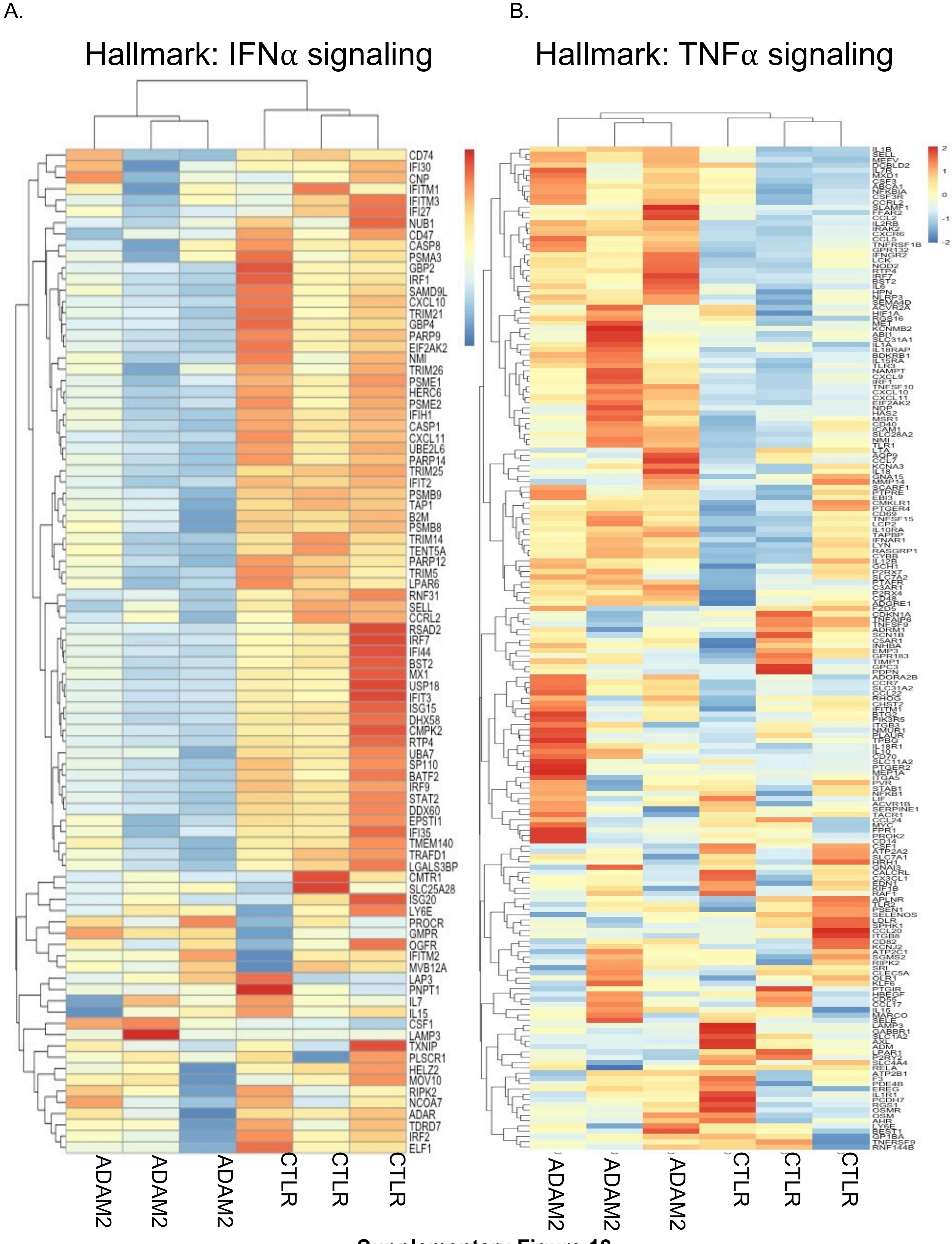


**Supplementary Figure 18**

A. Hallmark: IL2_STAT5_signaling

B. Hallmark: IL6_JAK_STAT3 signaling

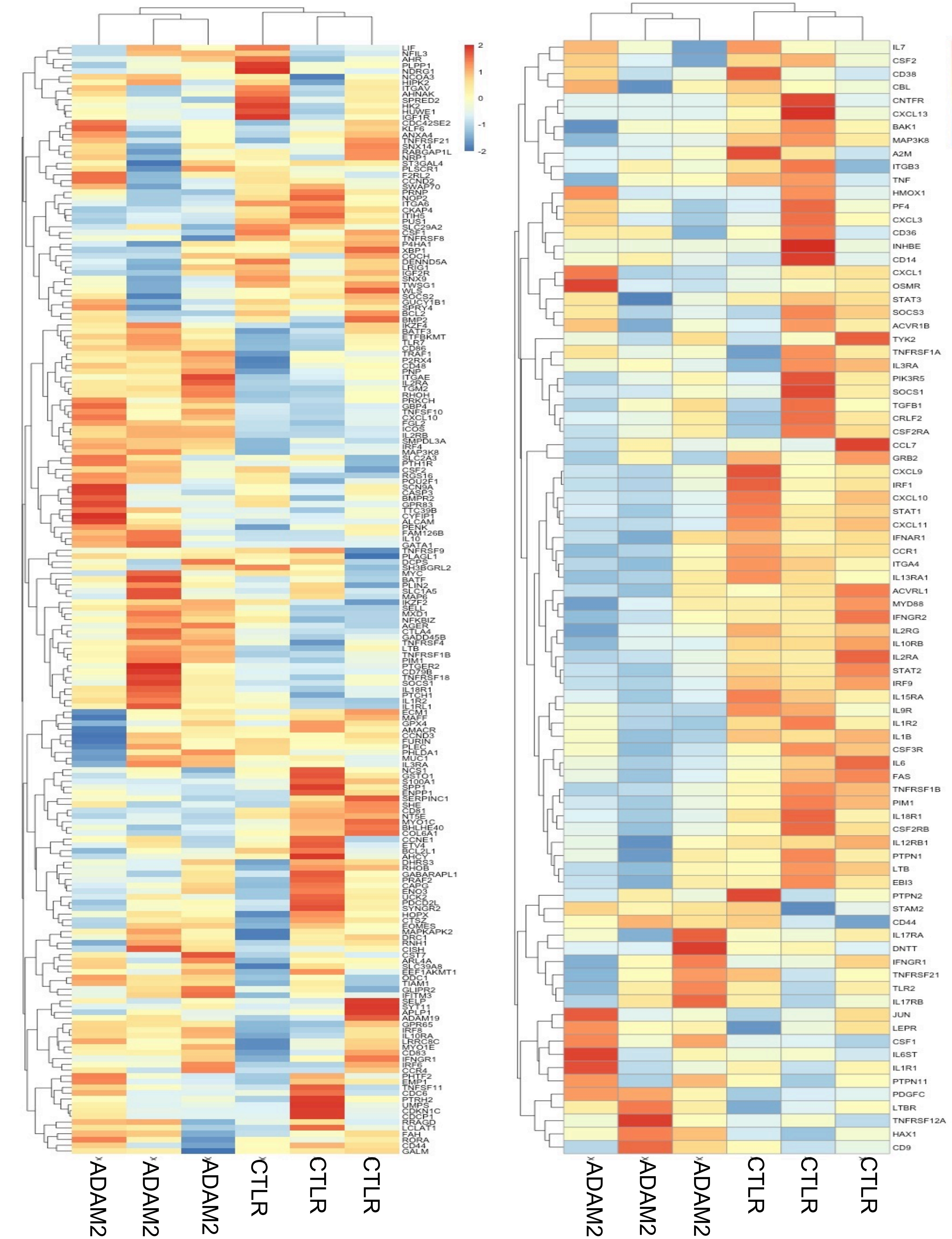


**Supplementary Figure 19**

A.

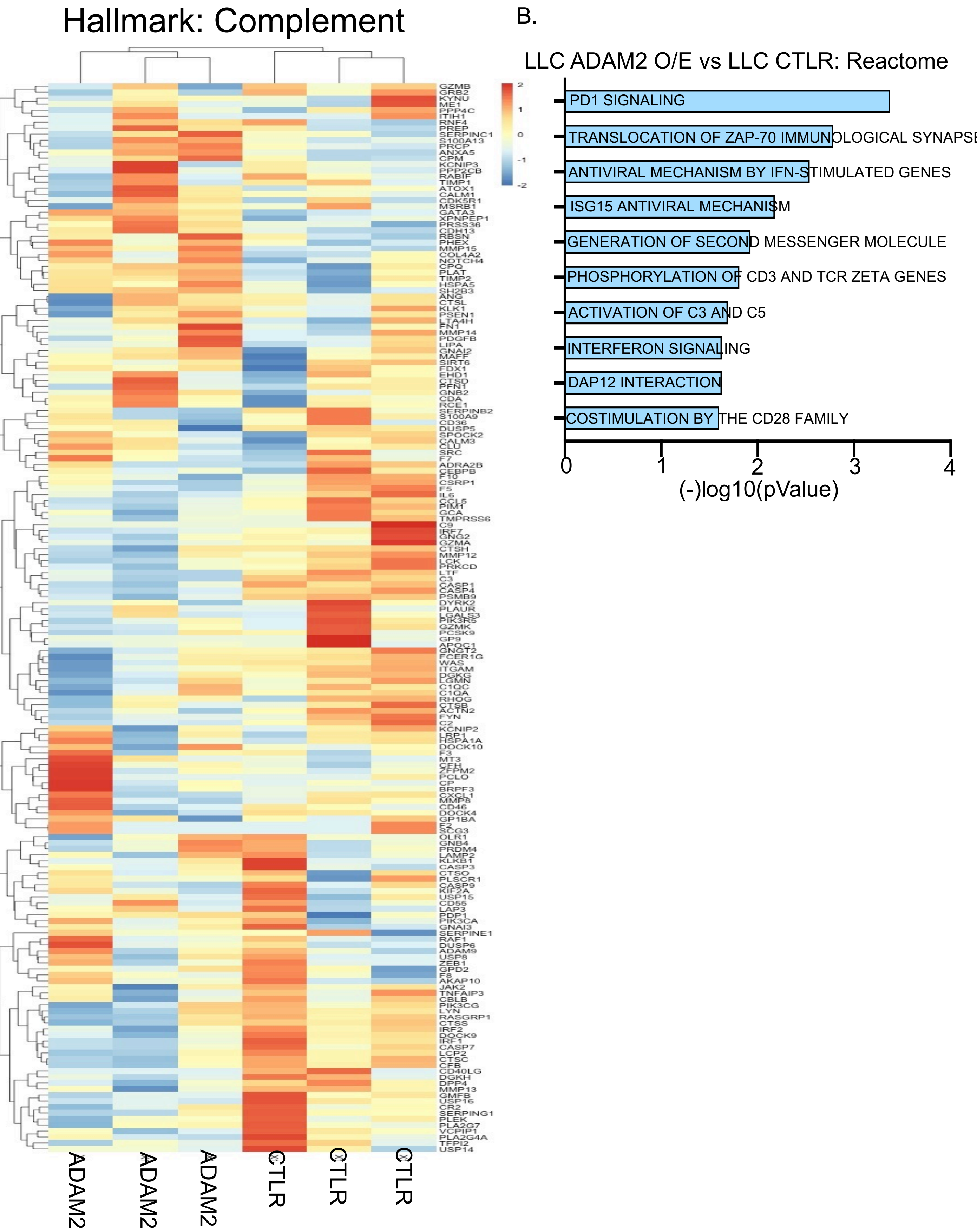


**Supplementary Figure 20**

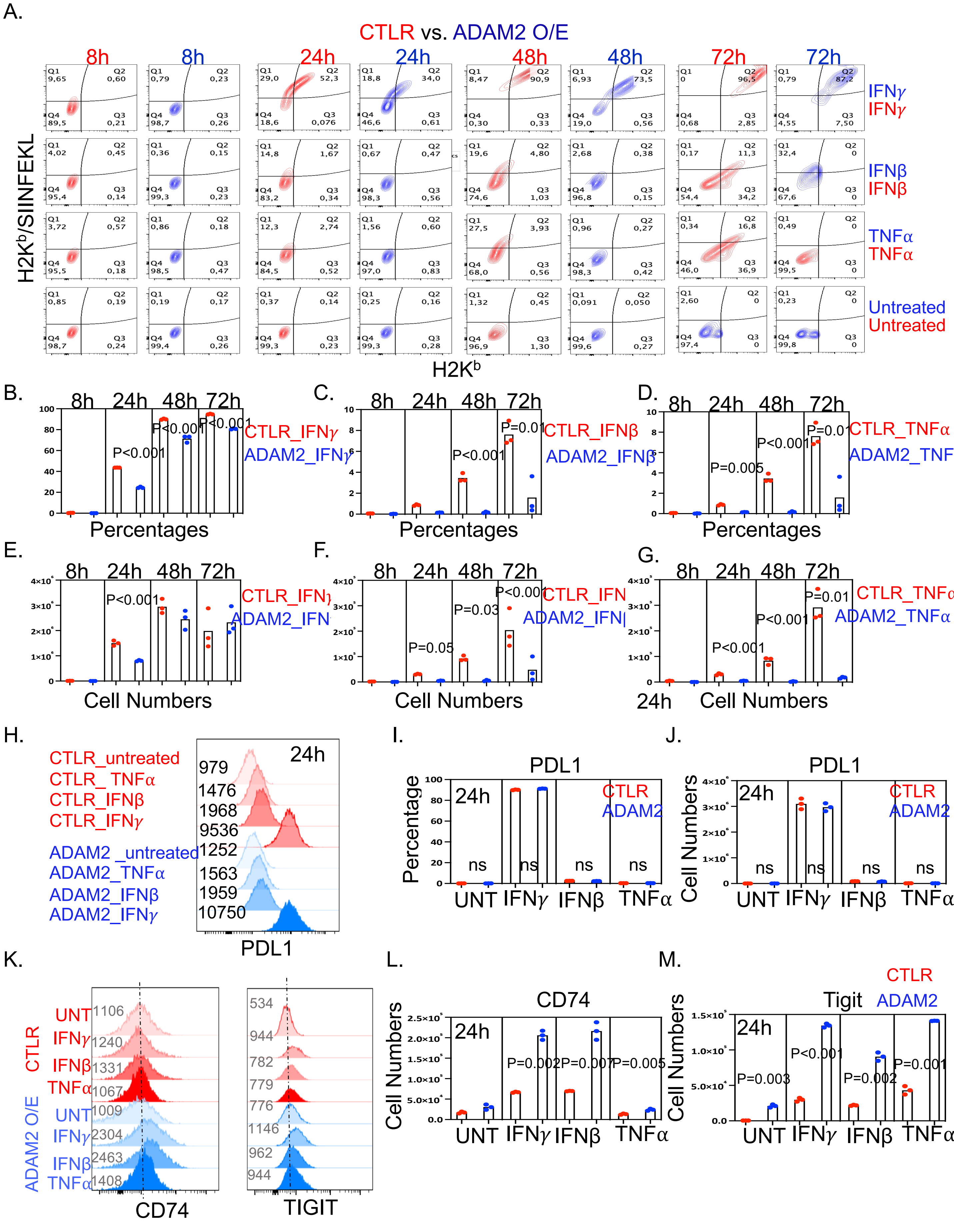


**Supplementary Figure 21**

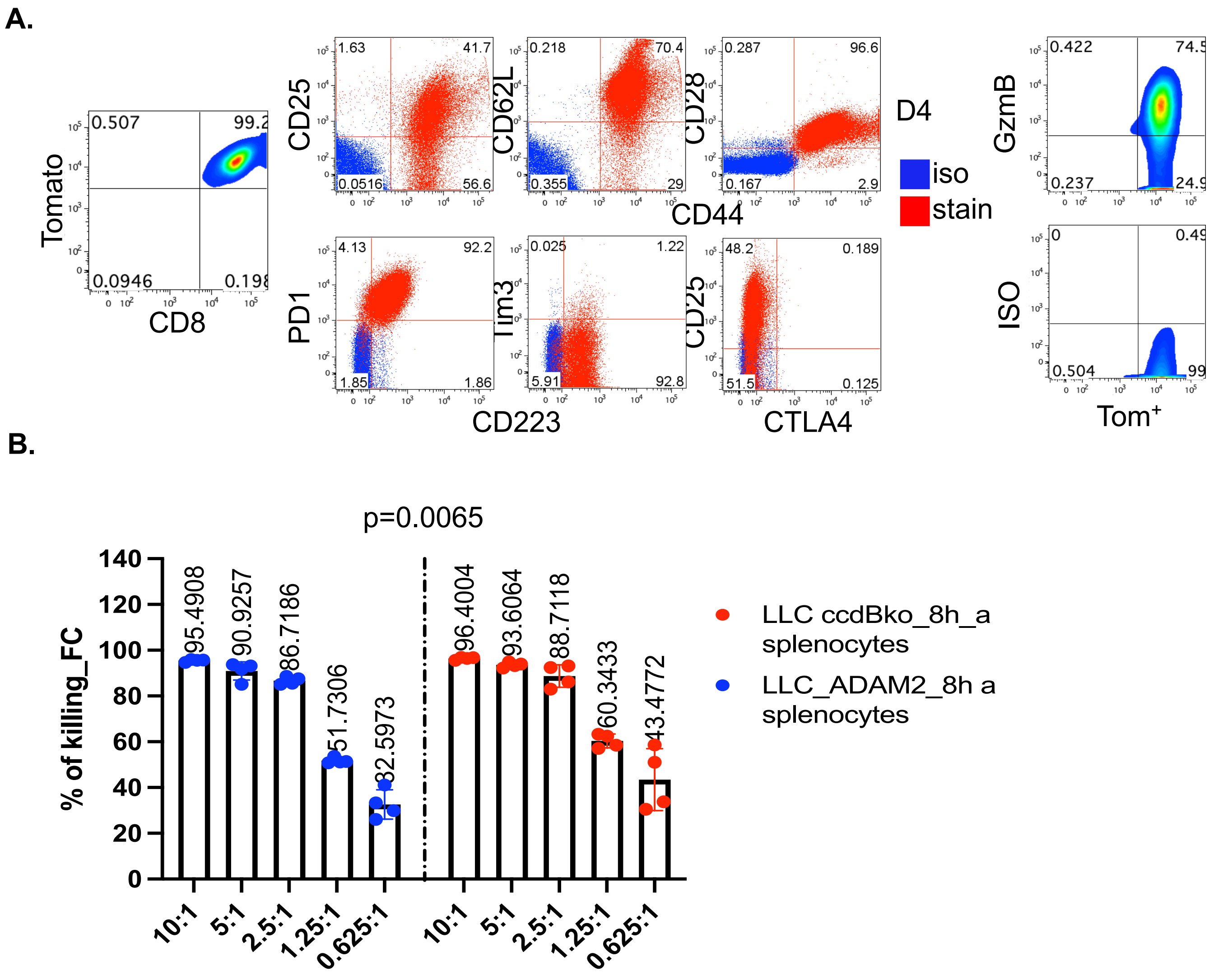


**Supplementary Figure 22**

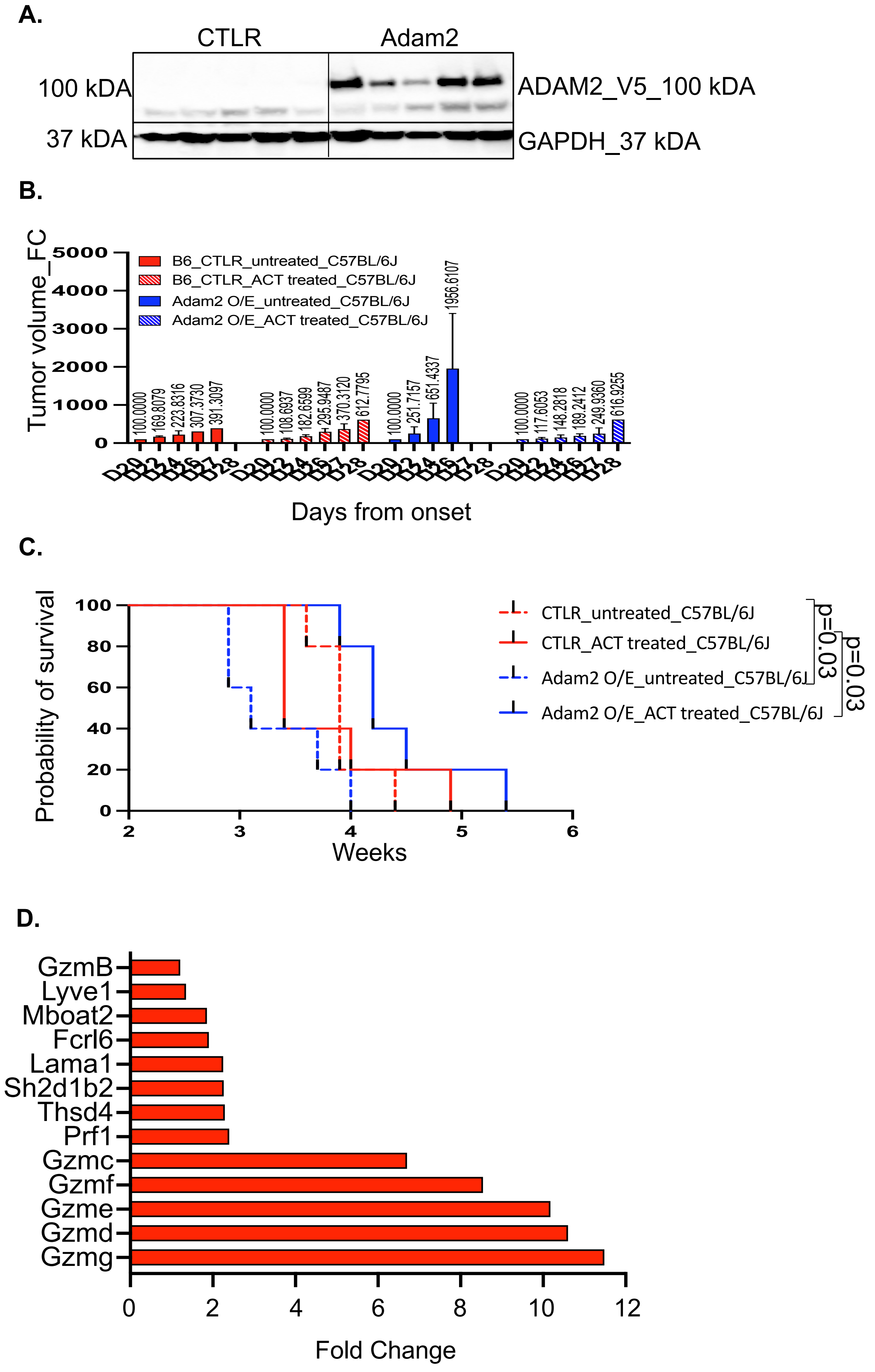


**Supplementary Figure 23**
